# Characterizing Bacterial and Archaeal Microbiomes of Singapore’s Urban Long-Tailed Macaques

**DOI:** 10.1101/2025.07.03.662964

**Authors:** Chissa-Louise Rivaldi, Justin J. S. Wilcox, Bailee Egan, Benjamin Gombash, Hope Hollocher

## Abstract

Animal microbiomes are sources of ecological responsiveness in the face of environmental change, and may serve as modulators and indicators of adaptability and stress in the face of emerging ecological perturbations. Baseline characterizations of non-human primate microbiomes will be important to applied and theoretical applications of microbiome research. Long-tailed macaques (*Macaca fasciularis*) are among the most ubiquitous primates, they live in close association with humans, and are a common model organism in biomedical research. Here, we use 16S amplicon metabarcoding of the V4 hypervariable region to provide baseline information on taxonomic and inferred functional variation in the prokaryotic (Archaea and Eubacteria) assemblies of oral and gut microbiomes of urban long-tailed macaques in Singapore. Oral microbiomes showed the most pronounced community structure at lower taxonomic levels, particularly ASVs. Gut microbiomes, in contrast, showed the most pronounced community structure at higher taxonomic levels, particularly phyla. Gut microbiomes showed clear groupings based on relative abundances of Proteobacteria, Firmicutes, and Bacteroidetes. Gut microbiome community composition was almost entirely explained by the Proteobacteria:Firmicutes ratio and this metric explained twice as much inferred functional variation as the more traditional Firmicutes:Bacteroidetes ratio. Archaeal communities in both oral and gut communities were dominated by methanogens. These were the only archaea found in the gut, but ammonia-oxidizing archaea were consistent constituents of oral microbiome assemblies as well. Ultimately, our findings imply distinct microbiome composition in Singapore’s urban macaques relative to reports from non-urban conspecifics.

## INTRODUCTION

Animals possess complex and variable microbiomes that are critical to their physiology and ecological responsiveness. Medical research has demonstrated clear and important impacts of these microbial communities on human phenotypes, such as obesity (Turnbaugh et al., 2006), gastrointestinal distress (Sartor, 2008), and nervous system function (De Vadder et al., 2018). Microbiomes certainly play an important role in the physiology and ecology of other metazoans as well (Hammer et al., 2019; Lee & Hase, 2014), and efforts to classify the microbiomes of non-human species are ongoing (Levin et al., 2021). However, the extent to which microbial communities converge and diverge in taxonomy and function across hosts remains unresolved (Bornbusch et al., 2023; D. Li et al., 2024; Song et al., 2019). As such, cross-species comparisons of microbiome composition and ecology depend on the identification of consistent phylogenetic and ecological influences and constraints (Mallott & Amato, 2021). Similarities between humans and non-human primates (NHPs) make the latter promising candidates for the expansion of microbiome research into non-model organisms (Amato, 2013; Clayton et al., 2018; Sharma et al., 2021).

NHPs are also implicated in a range of ongoing ecological challenges, including climate change (Bernard & Marshall, 2020), loss of habitat (Fernández et al., 2022), zoonotic disease transmission (Burgos-Rodriguez, 2011; Davoust et al., 2018), and reverse-zoonoses resulting from increased exposure to humans (Noman et al., 2024; Weary et al., 2024; Wilcox et al., 2021). Patterns of human land use are central to all of these threats (Fernández et al., 2022) and directly influence risk of bidirectional disease transmission between humans and other primates (Esposito et al., 2023; White & Razgour, 2020), including increased risk of major zoonotic and reverse-zoonotic events, and the emergence of new infectious diseases in human and non-human primates alike. Such threats are likely to be particularly pronounced where there are major losses of biodiversity, major increases in human population density, and increased interactions between human and non-human primates. Extensive human development—such as urbanization—may therefore exacerbate many of the challenges facing non-human primates.

Microbiomes may present a source of plasticity with which non-human primates may adapt to these new challenges (Lynch & Hsiao, 2019; Stumpf et al., 2016; West et al., 2019). They may also act as indicators of disturbance to inform conservation efforts (Stumpf et al., 2016; West et al., 2019). Understanding the normal microbiota and their variability within non-human primates is essential to theoretical projections and conservation applications of microbiome research to NHPs as well as identifying and differentiating zoonotic and reverse-zoonotic disease risk that may arise from interactions between human and non-human primates (Patouillat et al., 2024).

Baseline characterizations of microbiomes have been published for many primates (Clayton et al., 2018). Bacterial communities of primates are highly variable and responsive to changes in ecological conditions. The Firmicutes:Bacteroidetes (F:B) ratio is commonly used to assess changes in bacterial communities within humans (Ross et al., 2024) and other primates (Barelli et al., 2020; Kalkeri et al., 2021; Sawaswong et al., 2021), but the breadth of taxa involved in ecological responsiveness and their consistency of functions are poorly resolved across primates (Amato et al., 2015; Bornbusch et al., 2023; Clayton et al., 2018; Wills et al., 2022). Symbiotic archaea of NHPs are not well characterized (Manara et al., 2019), but archaea tend to perform unique and mutualistic functions in other hosts (Weiland-Bräuer, 2023). This may make archaea an important and neglected source of adaptability within NHPs. Archaea are also the predominant source of biological methane production (methanogenesis) on Earth, making shifts in archaeal abundances in response to changing ecological conditions of significance to global carbon cycles and climate change (Lyu et al., 2018; Moissl-Eichinger et al., 2018). Bacterial communities of humans show predictable shifts in response to collinear urbanization, industrialization, and consumption of ultra-processed foods (Mancabelli et al., 2017; Sonnenburg & Sonnenburg, 2019). Studies on urban foraging NHPs are lacking, but may help to elucidate conserved or divergent patterns of microbiome responses to industrial impacts across primates more generally and reveal new pathological states associated with industrialization (Grant et al., 2019). The archaeal and bacterial communities of non-human primates in urban settings are therefore of broad interest to questions of microbiome function, conservation, and health.

Long-tailed macaques (*Macaca fascicularis*) are ubiquitous NHPs of particular interest as a model organism in medical studies (Guebre-Xabier et al., 2020; Uno et al., 2016), human-associated edge species (Gumert, 2011), and potential sources of zoonotic infections (Kaewchot et al., 2022; M. I. Li et al., 2021). The macaques of Singapore live in a highly urbanized environment where they frequently associate with humans. Raiding of anthropogenic food sources is common, although the degree of anthropogenic impact varies between troop ranges (Klegarth, Hollocher, et al., 2017). As such, Singapore’s macaques are excellent targets for studies on the impacts of urbanization on NHPs (Fuentes et al., 2008; Gumert, 2011). A recent study on the microbiomes of captured and wild long-tailed macaques in Thailand shows clear distinctions in both the oral and gut microbiomes of these groups and links these to changes in the F:B ratio (Sawaswong et al., 2021). As this study was conducted in non-urban settings and did not characterize archaea, characterizations of urban bacterial and archaeal communities of these macaques are therefore lacking.

Here, we describe the community structure and dominant taxa of the oral and gut prokaryotic microbiomes of long-tailed macaques in Singapore. Using the V4 region of the 16S rRNA gene, we assess the overall bacterial and archaeal composition of body sites (oral and gut), identify which taxa define body sites, and measure differences in community structure (richness, evenness, dispersion, and dissimilarity), in order to provide baseline metrics of these bacterial communities in urban macaques. We inspect these communities at the phylum, family, and amplicon sequence variant (ASV) levels to identify the taxonomic levels at which these communities are structured and possible drivers of these changes. We first aim to answer the question of how the oral and gut microbial communities of macaques living in Singapore are structured by body site. We then evaluate the taxonomic levels and taxa that structure these communities within body sites. After identifying Proteobacteria as a unique source of variation within these communities, we compare the relative efficacy of the Proteobacteria:Firmicutes (P:F) ratio to the more traditional F:B ratio to assess whether the inferred functional profiles of macaque microbiomes mirror those of humans or follow their own trajectories.

## MATERIALS AND METHODS

### Study Site, Sampling, and DNA extraction

Saliva and fecal samples (N=92, 46 saliva and 46 fecal) were collected from free-ranging long-tailed macaques across ten sites throughout Singapore (Figure 1; Supplementary Table S1). Fecal (five sites, N=46) and saliva (nine sites, N=46) samples were collected from Singapore in June 2013 – October 2013. Collection times fall within Southwest Monsoon season, marked by frequent rain (Meteorological Service Singapore, n.d.). Fresh fecal samples were recovered from the ground when defecation was observed; all fecal samples from each site were collected on a single day and time (usually the evening) to prevent multiple sampling of the same individual. Fecal samples were kept on dry ice during the day of collection in the field and then stored at - 85C. All samples were also shipped on dry ice and stored in a −85C freezer until final processing (Klegarth et al., 2017; Wilcox and Hollocher, 2018). Saliva samples were collected using sugar-syrup flavored buccal swabs from Salimetrics, offered to the macaques and recovered immediately after discard when only a single individual had handled it (Klegarth, Sanders, et al., 2017). Swabs were kept on ice packs during collection and stored in 1% sucrose cell lysis buffer at −85°C (Wilbur at al., 2012). Under these collection conditions ground contamination and delay in processing should be insufficient to affect downstream analyses (Grieneisen et al., 2019). Total DNA was extracted from feces using the Qiagen QIAMP Stool Minikit (Qiagen, Hilden, Germany) as per manufacturer instructions (Qiagen, 2012). DNA from saliva samples was extracted via a phenol chloroform protocol: samples were incubated overnight at 55°C in 900.00 µL CTAB with 20.0 µL Proteinase K enzyme. Samples were shaken with phenol/chloroform for one minute, and then spun to separate the aqueous phase that was then washed with 100% EtOH, followed by washing with 70% EtOH solution, and stored at −80°C. Further details regarding sample collection and DNA extraction are described previously (Klegarth, Sanders, et al., 2017; Wilcox & Hollocher, 2018).

**Figure 1.**
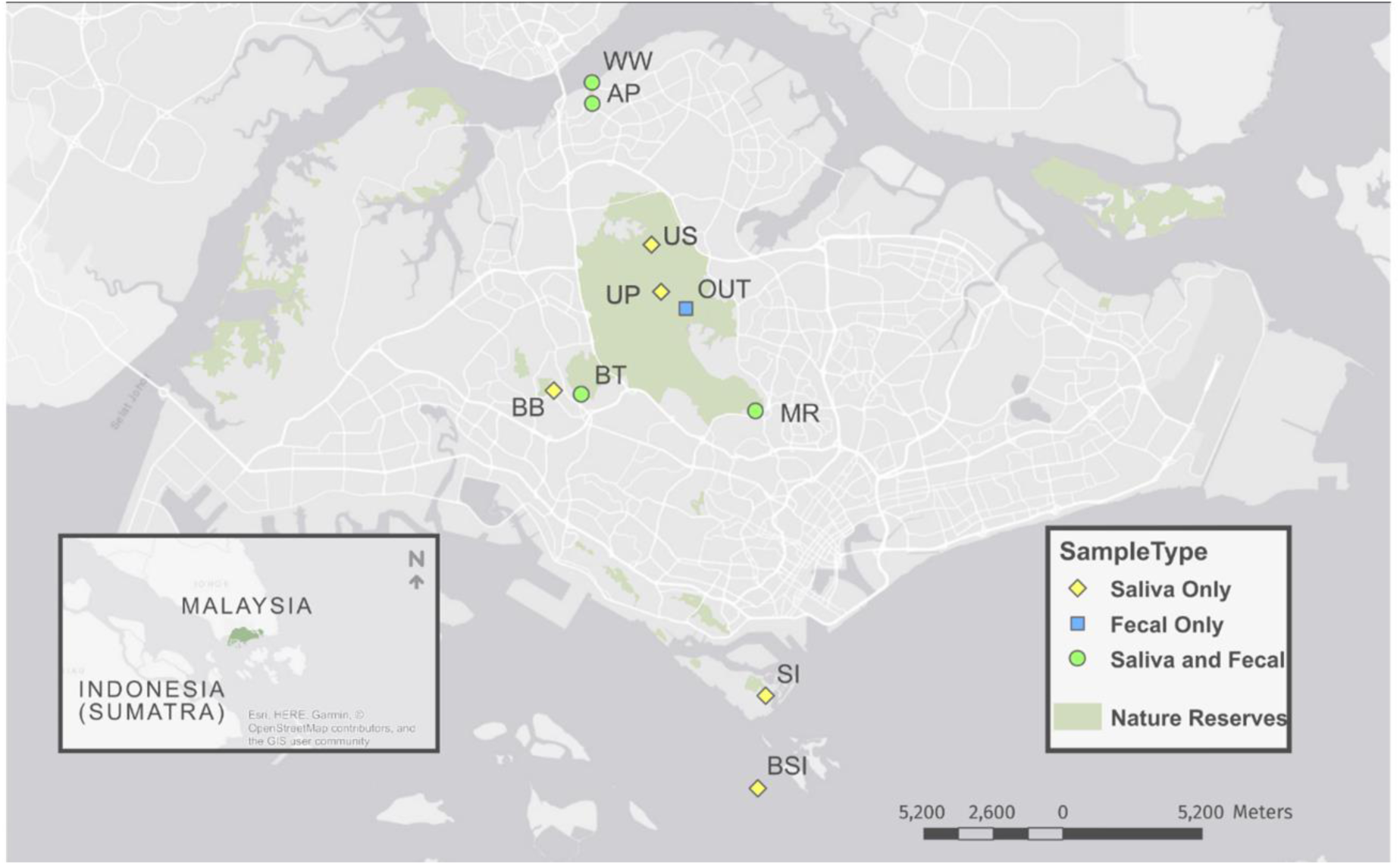
Sampling sites indicate populations from which samples were obtained. Samples were collected throughout accessible macaque habitat. Geographic coordinates and number of samples for each site are as follows: AP - Admiralty Park: 1° 27’ 2.9844’’ N, 103° 46’ 38.9964’’ E (N = 12); BB - Bukit Batok: 1° 22’ 36.2900” N, 103° 45’ 30.6242” E (N = 3); BSI - Big Sister’s Island: 1° 12’ 52.4916” N, 103° 50’ 6.3055” E (N = 8); BT - Bukit Timah: 1° 22’ 47.0205” N, 103° 46’ 25.42636” E (N = 11); MR - MacRitchie Reserve 1° 20’ 28.0572’’ N, 103° 49’ 48.5616’’ E (N = 17); OUT - Old Upper Thomson Road: 1° 22’ 49.7233” N, 103° 49’ 6.3001” E (N = 15); SI - Sentosa Island: 1° 15’ 29.3885” N, 103° 48’ 33.5866” E (N = 4); UP - Upper Pierce: 1° 22’ 32.2896” N, 103° 48’ 17.3988” E (N = 6); US - Upper Seletar: 1° 23’ 55.5432” N, 103° 48’ 23.2812 E; WW - Woodlands Waterfront: 1° 27’ 6.6652’’ N, 103° 46’ 52.2617’’ E (N = 11).

All research conducted within this study complied with protocols approved by the University of Notre Dame’s Institutional Animal Care and Use Committees (IACUC; protocol IDs 07-001, 14-002, and 14-05-1835) and also adhered to the American Society of Primatologists Principles for the Ethical Treatment of Non-Human Primates. Samples were collected with permission from the Singapore National Parks Board (permit #NP/RP11-029) and exported under CITES (11SG006180CE, 12SG006486CE, 14SG009020CE) and CDC protocol numbers (2006-10-148, 2011-01-111, 2012-06-034, 2014-10-096).

### Target Amplification and High-Throughput Sequencing

A 287 bp sequence of the V4 region of the 16S rRNA gene was amplified from genomic DNA using the following primers (Illumina adaptors in italics): Forward (S-D-Arch-0519-a-S-15, 5’ - *TCG TCG GCA GCG TCA GAT GTG TAT AAG AGA CAG* CAG CMG CCG CGG TAA −3’), Reverse (S-D-Bact-0785-b-A-18, 5’ - *GTC TCG TGG GCT CGG AGA TGT GTA TAA GAG* ACA GTA CNV GGG TAT CTA ATC C −3’) (Klindworth et al., 2013; Van Bleijswijk et al., 2015) in the following PCR conditions: for a 25.0 µL reaction volume, 10.00 µL ddH2O, 12.50 µL Kappa HiFi Hot Start Taq Ready-Mix (Kappa Biosystems, Wilmington, MA, USA), 0.75 µL 10 µM forward and reverse primer, 1.00 µL template DNA. Thermocycler conditions were: 3:00 initial denaturation at 95.0 °C, 30 cycles (0:20 denaturation at 95 °C, 0:15 anneal at 65 °C, 0:15 extension at 72 °C), and a final extension time of 3:00 at 72 °C. Included in the first and last wells of the 96-well plate were negative controls, containing only autoclaved water in the place of the template DNA. Next to these negative controls were reactions containing genomic DNA from a prepared mock community (ZymoBIOMICS TM Microbial Community DNA Standard, Catalog no: D6306) as template, included to identify potential bias from PCR and sequencing assays and improve reproducibility in our work.

### Illumina PCR and sequencing

PCR products were purified with Ampure XP Beads (Beckman Coulter, Indianapolis, IN, USA), and size distributions were taken from each library using a Bioanalyzer DNA 7500 chip (Agilent, Santa Clara, CA, USA). Size distributions were unique to each library, and no major anomalies were seen. Index PCRs were performed on all libraries following clean-up and quality assessment using a 95 °C denaturing phase, 55 °C annealing phase, and 72 °C extension phase, for eight cycles, each lasting 30 seconds.

Libraries were pooled and normalized prior to index amplification. Each library was used to create 100 µM aliquots based on the following equation:

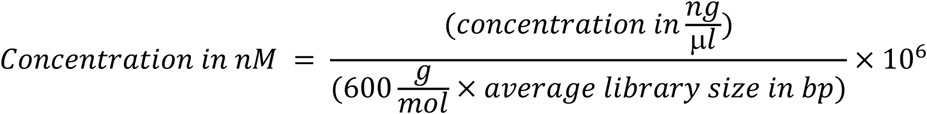

A normalized library was prepared by diluting 5.0 µL from the prepared library to 20.0 nM aliquots. Size distributions were bioanalyzed using a DNA 7500 chip, to confirm appropriate size distributions within the library. The library was sequenced on an Illumina HiSeq 2500 (Illumina, San Diego, CA, USA), using a rapid run with paired-end 250 bp reads at the University of New Hampshire Hubbard Center for Genome Studies.

### Bioinformatic Workflow and Statistical Analysis

The R package Dada2 was used for processing sequences, including filtering, trimming, and merging, and removal of chimeras (Callahan et al., 2016). Taxonomy assignment was performed using the Dada2 function of the RDP classifier and the RDP training set (release 11.5), and all eukaryotes and Cyanobacteria were removed (Callahan et al., 2016). All ASVs were retained hereafter. Rarefaction curves were created with the rarecurve function from the vegan package (v 2.5-7(Oksanen et al., 2022). We aligned the contents of mock community samples to a reference database of 16S ssuRNA sequences downloaded from the manufacturers website (zymoresearch.com, version 2) using HISAT2 (v. 2.1.0; (Kim et al., 2015) with greater than 98% of reads aligning to the reference. Negative controls had 2142 and 2406 reads total, representing 0.14% (lowest control as proportion of highest sample) to 1.23% (highest control as proportion of lowest) of the read depths of sequenced samples. Based on these findings samples were considered to be sufficiently pure that no corrections were made for technical contamination.

Microbiome α-diversity (Shannon), richness and Pielou’s evenness were calculated in R using the vegan package. Rarefaction curves confirmed consistent richness in all samples well below sequencing depth; therefore, no other transformations were applied in the α-diversity metrics. To assess dispersion by body-site, Bray-Curtis dissimilarity was assessed for each group based on the spread of samples around the group’s centroid. To account for uneven sampling, we used Mann-Whitney U-tests to determine differences in richness, evenness, and dissimilarity between body sites.

All distance-based statistics used relative abundances. Relative abundances were preferred to more complex methods of normalization as they have been suggested to introduce the fewest false positive artifacts into analyses (McKnight et al., 2019). To inspect taxa defining the oral and gut environments, we employed a K-means clustering method, and identified the taxa correlated with each cluster. A K-value of two was selected in an attempt to show that the oral and gut bacterial communities are distinctive. If these two clusters match the sample type, then it would support body site distinctiveness. The adjusted Rand index was used to measure agreement between the clusters and sample type (Hubert & Arabie, 1985). Euclidean distances between samples were calculated using the vegdist function in the vegan package (v 2.5-7) in R. The envfit function in the vegan package was used to determine which bacterial taxa were strongly correlated (R^2^ > 0.7) with each cluster.

NMDS plots were created to visualize grouping in samples in R using the vegan package, using Bray-Curtis distances. One-way PERMANOVAs (adonis2 function from the vegan package in R) were used to test for differences in composition between body sites and geographical sampling locations within body sites at phylum, family levels, and ASV levels. We measured the dispersion of variance between body sites and then between sampling locations within body site using the betadisper and permutest functions in the vegan package. Principal component analyses for drivers of assembly composition were performed in base R using the prcomp function for ASVs and families—due to their high number of taxa to sample ratio—and the princomp function for phyla and graphed using the ggbiplot package (version 0.55). Changes in relative abundances of families within the phyla Proteobacteria, Firmicutes and Bacteroidetes, were tested using a series of Mann-Whitney U tests as a post-hoc—Dunn–Šidák corrected α-values were used to control for multiple tests. MetaCyC functional pathways were predicted using the software PICRUSt2 using all ASVs (Barbera et al., 2019; Caspi et al., 2014; Czech et al., 2020; Douglas et al., 2020; Louca & Doebeli, 2018; Mirarab et al., 2011; Ye & Doak, 2009). The resulting pathway table was then analyzed in R using ALDEx2, which uses a Dirichlet-multinomial model to infer abundance from counts, to determine which pathways were significantly associated with body sites (v 1.22.0) (Fernandes et al., 2013, 2014; Gloor et al., 2016). As recommended by the authors of this package, we used an effect size of > 1.0 to determine significance of inferred pathways associated with body sites. All scripts have been made available as supplementary code.

## RESULTS

A total of 71,460,186 paired-end reads were obtained for processing. After filtering and removal of chimeras, eukaryotes, and Cyanobacteria, 52,192,561 reads remained. We detect two domains (Archaea and Bacteria), 33 phyla, 237 families, and 14,892 ASVs across all body sites and samples. ASV counts plateau in rarefaction analysis before maximum reads depths are reached in all samples (Supplementary Figure 1).

The five most abundant phyla in the sequencing data belonged to Bacteria (Figure 2) and accounted for greater than 97.9% of filtered reads. Bacteria consisted of Proteobacteria (39.4%), Firmicutes (34.1%), Bacteroidetes (12.3%), and Actinobacteria (7.8%), and Fusobacteria (4.4%). The five most abundant bacterial families accounted for 50.7% of reads and were all members of either Proteobacteria or Firmicutes: Enterobacteriaceae (13.3%, Proteobacteria), Streptococcaceae (12.3%, Firmicutes), Pasteurellaceae (9.7%, Proteobacteria), Moraxellaceae (9.3%, Proteobacteria), Planococcaceae (6.0%, Firmicutes). The five most abundant ASVs— ASVs 2 (6.2%), 1 (4.9%), 4 (4.4%), 3 (3.8%), and 5 (3.5%)—accounted for 23.0% of reads and belonged to the Genera *Escherichia*/*Shigella* (Family: Enterobacteriaceae; Phylum: Proteobacteria), Actinobacillus (Family: Pasteurellaceae; Proteobacteria), Enterobacter (Family: Enterobacteriaceae, Phylum: Proteobacteria), and Streptococcus (Family: Streptococcaceae; Phylum: Firmicutes—included in both ASV 3 and ASV 5), respectively.

**Figure 2.**
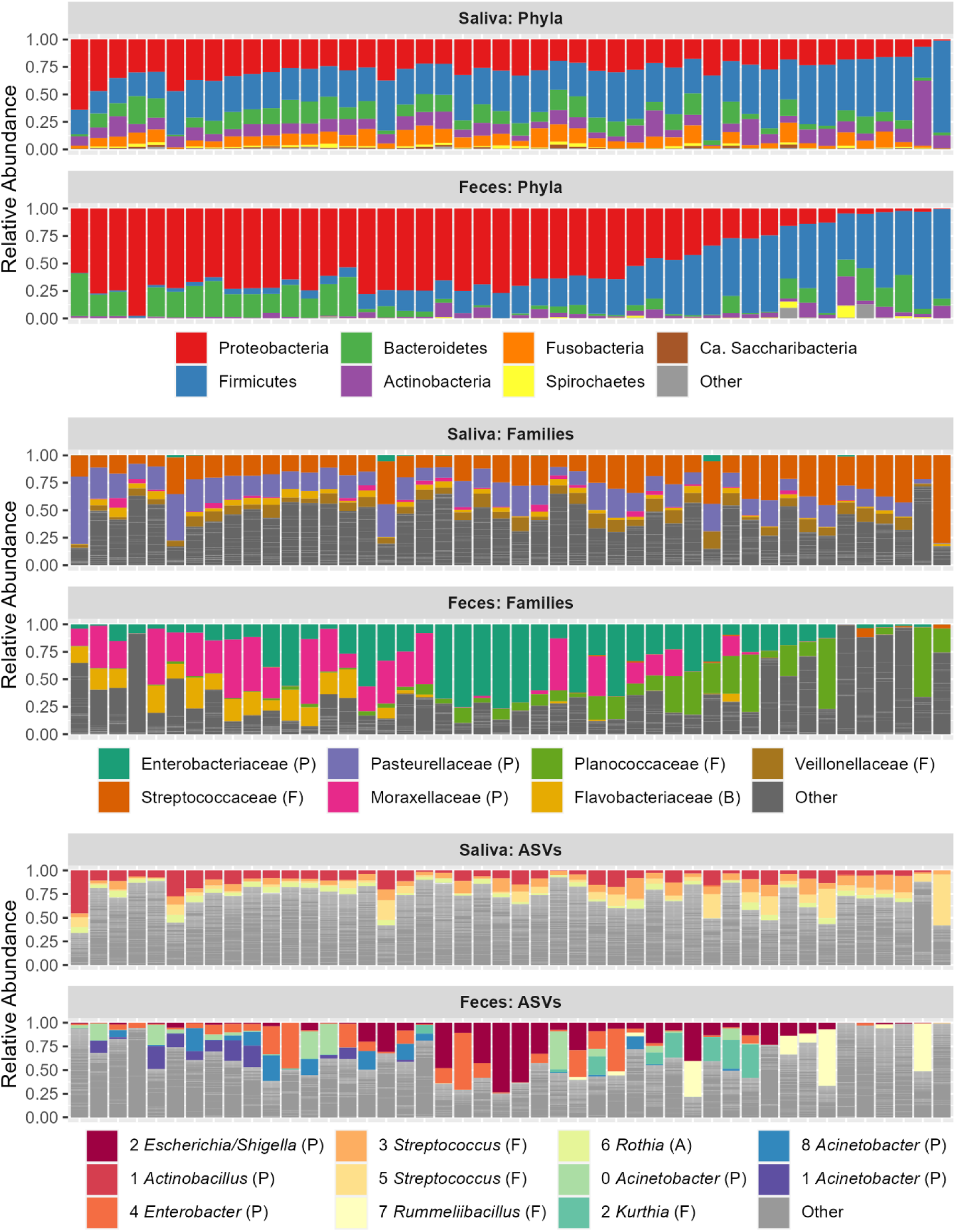
Relative abundance of bacterial phyla (top), Familes (middile), and ASVs (bottom) in saliva and fecal microbiomes. Samples are ordered left to right by decreasing ratio of Proteobacteria to Firmicutes (P:F) and colored by taxa. The most abundant taxa are shown with other taxa shown in gray; different groupings of other taxa are delineated by light gray lines.

Archaea accounted for less than 0.14% of reads. Five archaeal phyla were nonetheless represented (Supplementary Figure 2): Crenarchaeota (0.000014%), Euryarchaeota (0.13%), Pacearchaeota (0.000049%), Thaumarchaeota (0.000449%). Woesearchaeota (0.000008%).

Six archaeal families were found, consisting of Halobacteriaceae (0.000009%), Methanobacteriaceae (0.13%), Methanomassiliicoccaceae (0.006260%), Methanotrichaceae (0.000007%), Nitrosopumilus (0.000031%), and Nitrososphaera (0.000431%). In total, 43 archaeal ASVs were found of which only 21 could be assigned to a genus. Of these all were methane producing Euryarchaeota, with seven ASVs assigned to Methanobrevibacter and 10 assigned to Methanomassiliicoccus, one assigned to Methanosphaera, one assigned to Methanothrix, one to Methanobacterium, and one to Halalkalicoccus. No Nitrososphaera (Phylum: Thaumarchaeota) could be assigned to the genus level, but 13 ASVs of these presumably ammonia-oxidizing archaea were found.

### Describing the Oral Microbiome

The oral microbiomes contained two domains, 32 phyla, 213 families, and 10,339 ASVs. The five most common saliva phyla accounted for 97.1% of reads (Figure 2) and were Firmicutes (38.1%), Proteobacteria (27.0%), Bacteroidetes (11.8%), Actinobacteria (11.4%), Fusobacteria (8.8%). The five most abundant families accounted for 62.2% of reads with Streptococcaceae (24.2%; Firmicutes), Pasteurellaceae (19.3%; Proteobacteria), Veillonellaceae (7.7%; Firmicutes), Prevotellaceae (5.9%; Bacteroidetes), Micrococcaceae (5.0%; Actinobacteria). The five most abundant ASVs within saliva—1 (9.8%), 3 (7.6%), 5 (7.1%), 6 (4.1%), 9 (2.7 %)— accounted for 31.5% and belonged to the genera *Actinobacillus* (Proteobacteria), *Streptococcus* (Firmicutes), *Streptococcus*, *Rothia* (Actinobacteria), and *Streptococcus*, respectively. Archaea accounted for less than 0.001% of saliva reads, but contained five phyla, six families, and 38 ASVs. Euryarchaeota (0.009%) were dominant among these phyla followed by Thaumarchaeota (0.0009%), with the remaining phlyla showing relative abundances an order of magnitude or more below these. Most reads for saliva-dwelling archaea were assigned to methanogenic familes: Halobacteriaceae (0.000018%), Methanobacteriaceae (0.002692%), Methanomassiliicoccaceae (0.006176%), Methanotrichaceae (0.000013%). Two families of nitrogen-oxidizing archaea were also identified in saliva Nitrosopumilus (0.000062%) and Nitrososphaera (0.000861%). Six ASVs were assigned to the genera *Methanobrevibacter*. Eight ASVs belonged to *Methanomassiliicoccus*. One ASV belonged to *Halalkalicoccus*, and one belonged to *Methanothrix*.

### Describing the Gut Microbiome

The gut microbiomes contained two domains, 24 phyla, 165 families, and 5,480 ASVs. The five most abundant phyla accounted for more than 99.3% of reads and consisted of Proteobacteria (51.8%), Firmicutes (30.0%), Bacteroidetes (12.8%), Actinobacteria (4.1%), and Spirochaetes (0.6%). The five most abundant families in feces accounted for 67.6% of reads (Figure 2): Enterobacteriaceae (26.3%; Proteobacteria), Moraxellaceae (17.0%; Proteobacteria), Planococcaceae (12.0%; Firmicutes), Flavobacteriaceae (6.6%; Firmicutes), Lachnospiraceae (5.7%; Firmicutes). The five most abundant ASV—2 (12.6%), 4 (8.7%), 7 (4.2%), 10 (4.0%), and 12 (3.6%)—accounted for 33.1% of fecal reads. The first two of these belonged to the genera *Escherichia*/*Shigella* and *Enterobacter* and family Enterobacteriaceae, and the fourth to *Acinetobacter* within the family Moraxellaceae, all of which are Proteobacteria. Others belonged to the genera *Rummeliibacillus* and *Kurthia* within the family Planococcaceae (Phylum: Firmicutes), respectively. All reads attributed to archaea within feces belong to the phylum Euryarchaeota (0.2226797%). Archaea in feces were exclusively methanogenic, belonging to one of two families Methanobacteriaceae (0.259599454%) or Methanomassiliicoccaceae (0.006343091%) and the genera *Methanobrevibacter* (2 ASVs) and *Methanobacterium* (1 ASV) within the former and *Methanomassiliicoccus* (2 ASVs) within the latter.

### Comparing the Oral and Gut Microbiomes

K-means clustering with two *a priori* groupings did not allow for strong diagnostics of oral and gut microbiomes at ASV, family, or phylum level (Figure 3A), but performed best at the family level (adjusted Rand index value = 0.79217) and poorly at phylum level (adjusted Rand index value =0.36417) and ASV level (adjusted Rand index value = 0.14722). In contrast, unsupervised hierarchical clustering (UPGMA) of Bray-Curtis dissimilarities showed totally distinct groupings of samples by body site at the ASV, family, and phylum levels (Figure 3B). A PERMANOVA test of Bray-Curtis dissimilarities between body sites likewise revealed significant differences between clusters at the ASV (F=40.161, R^2^ =0.30855, p<0.0001), family (F=85.623, R^2^ =0.48754, p<0.0001), and phylum (F=30.645, R^2^ =0.25401, p<0.0001) levels, albeit with more variation within bodies sites than between them at all taxonomic levels (Figure 3C).

**Figure 3.**
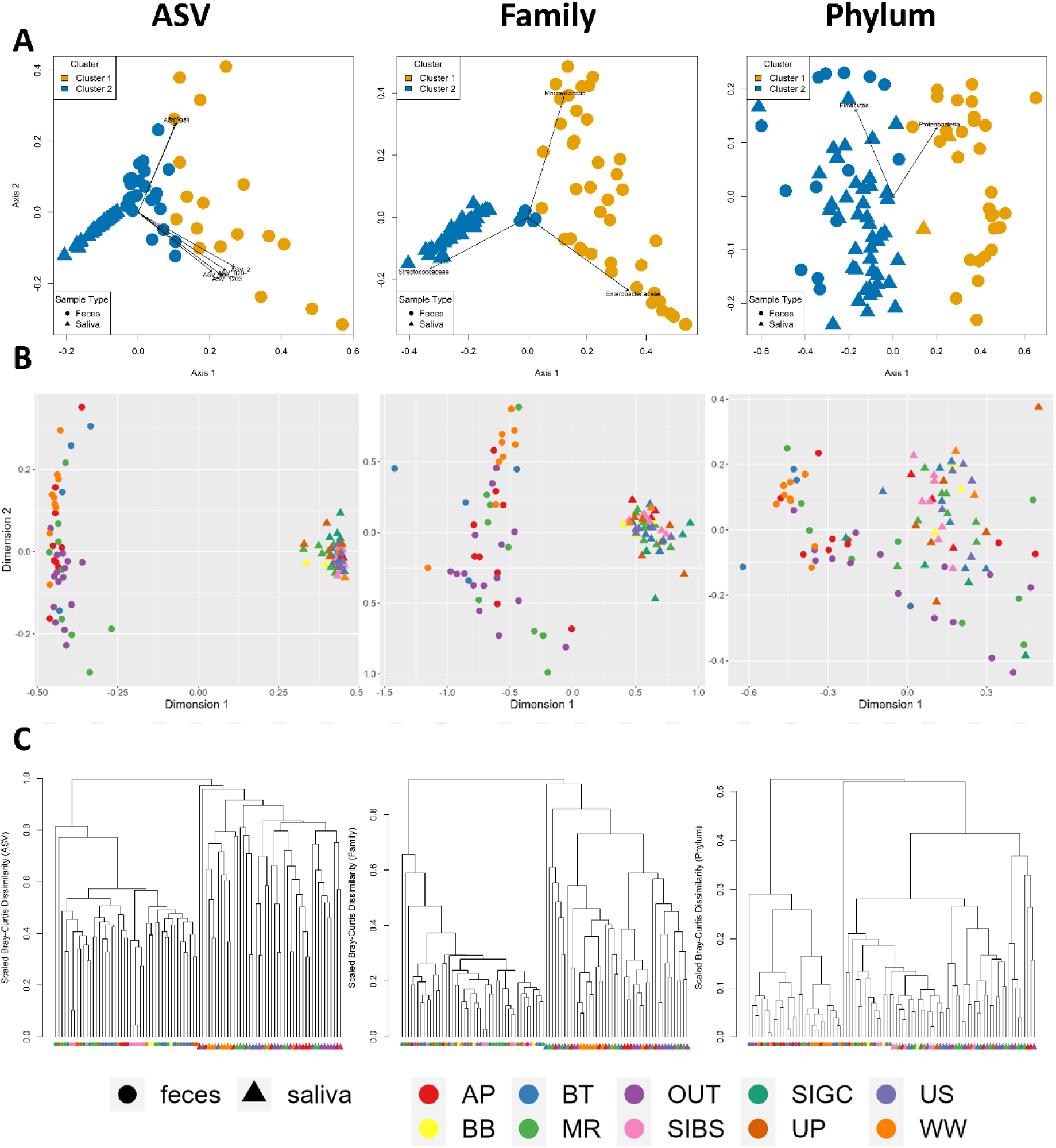
Clusters of oral and gut microbiomes using K-means clustering, Bray-Curtis MDS (B), and UPGMA clustering of Bray-Curtis dissimilarities (C). Taxonomic levels go from ASV to family to phylum (left to right). Body sites are shown by shapes in all figures. Assignment to K-means clusters are shown as colors in this analysis. Site locations are shown as colors for Bray-Curtis analyses. Stress on nMDS analyses are 7.496404, 10.08481, 8.662075 for ASVs, families, and phyla, respectively.

Taxonomic differences between oral and gut microbiomes were assessed using paired Mann-Whitney U tests on relative abundances with Dunn-Šidák corrections for false discovery rates. Analyses were conducted at the levels of ASVs, families, and phyla (Table 1). Four phyla— Deinococcus-Thermus, Fibrobacteres, Lentisphaerae, Proteobacteria—were enriched in fecal samples. Of these, the phylum Fibrobacteres occurred exclusively in the gut. In total, 15 phyla— notably, Acidobacteria, Actinobacteria, Spirochaetes, and a single archaeal phylum,

**Table 1.**
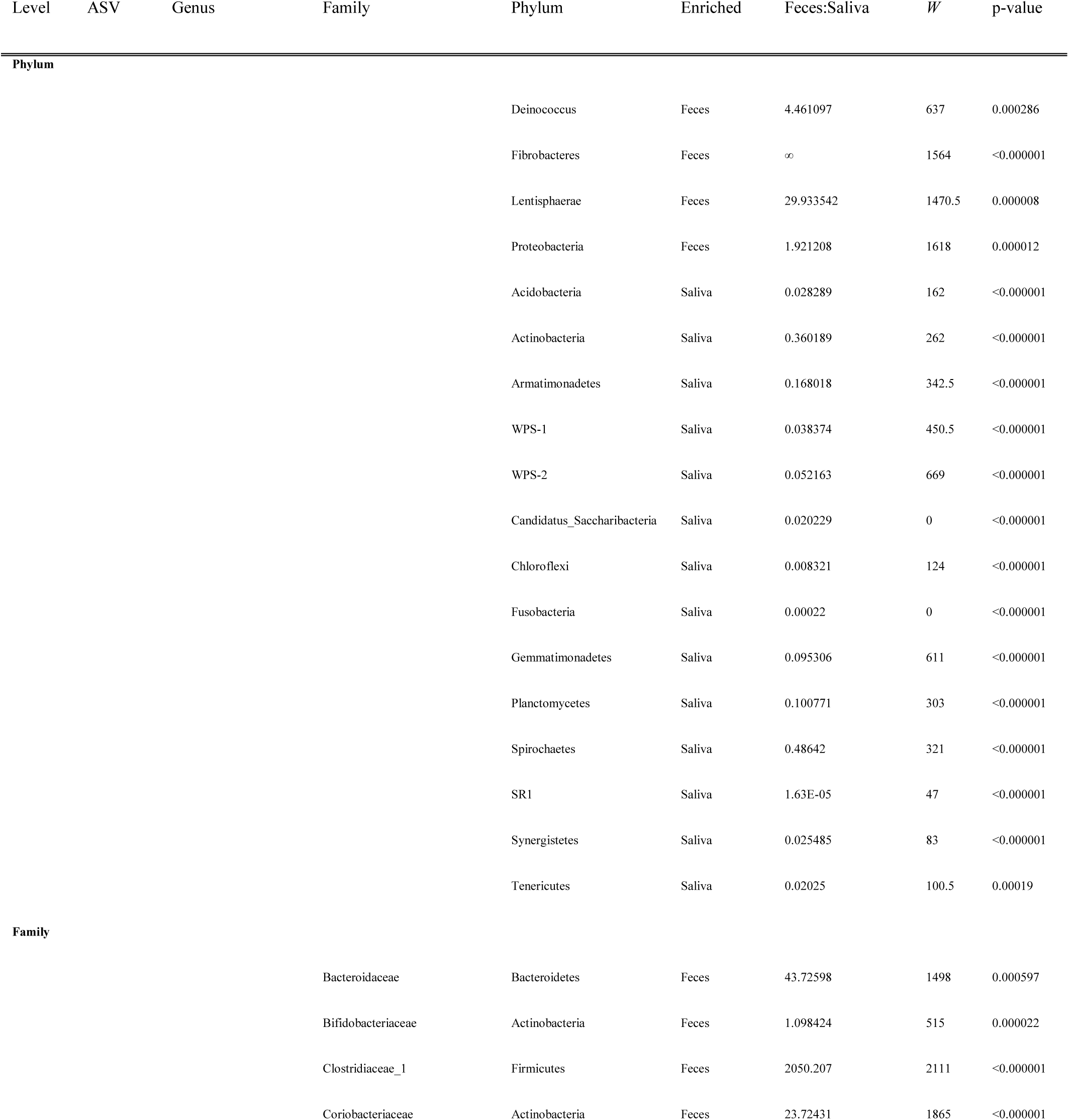
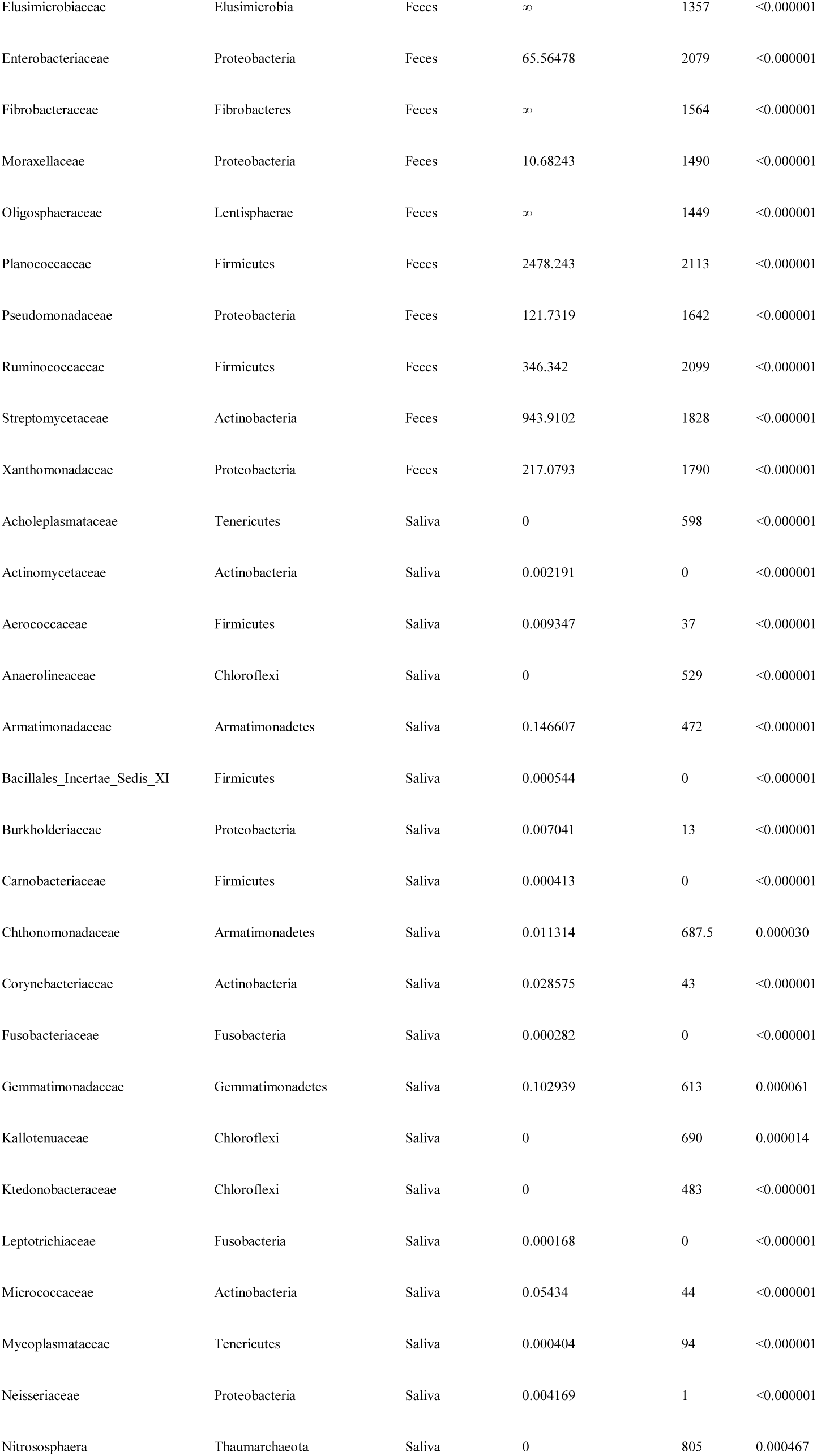
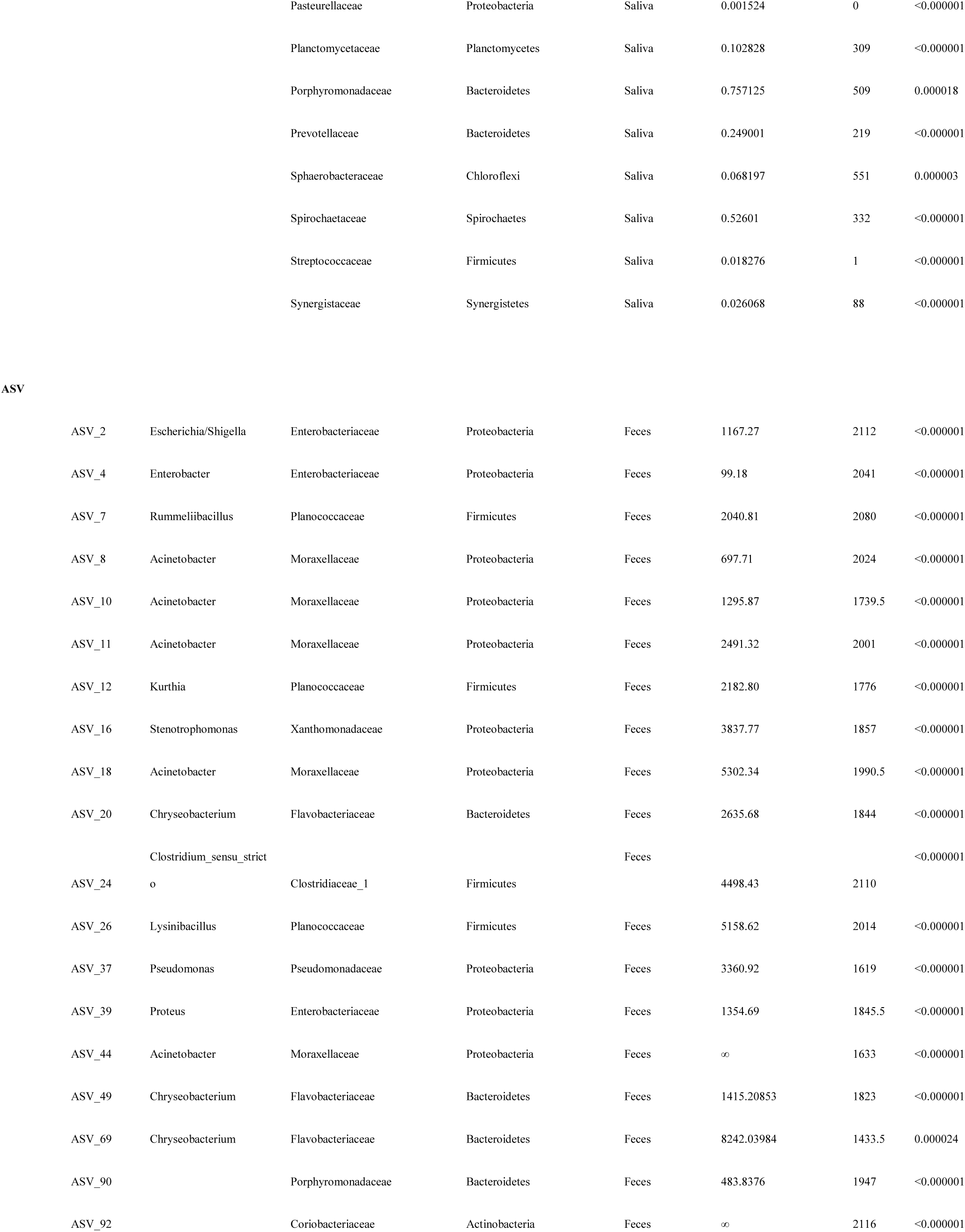
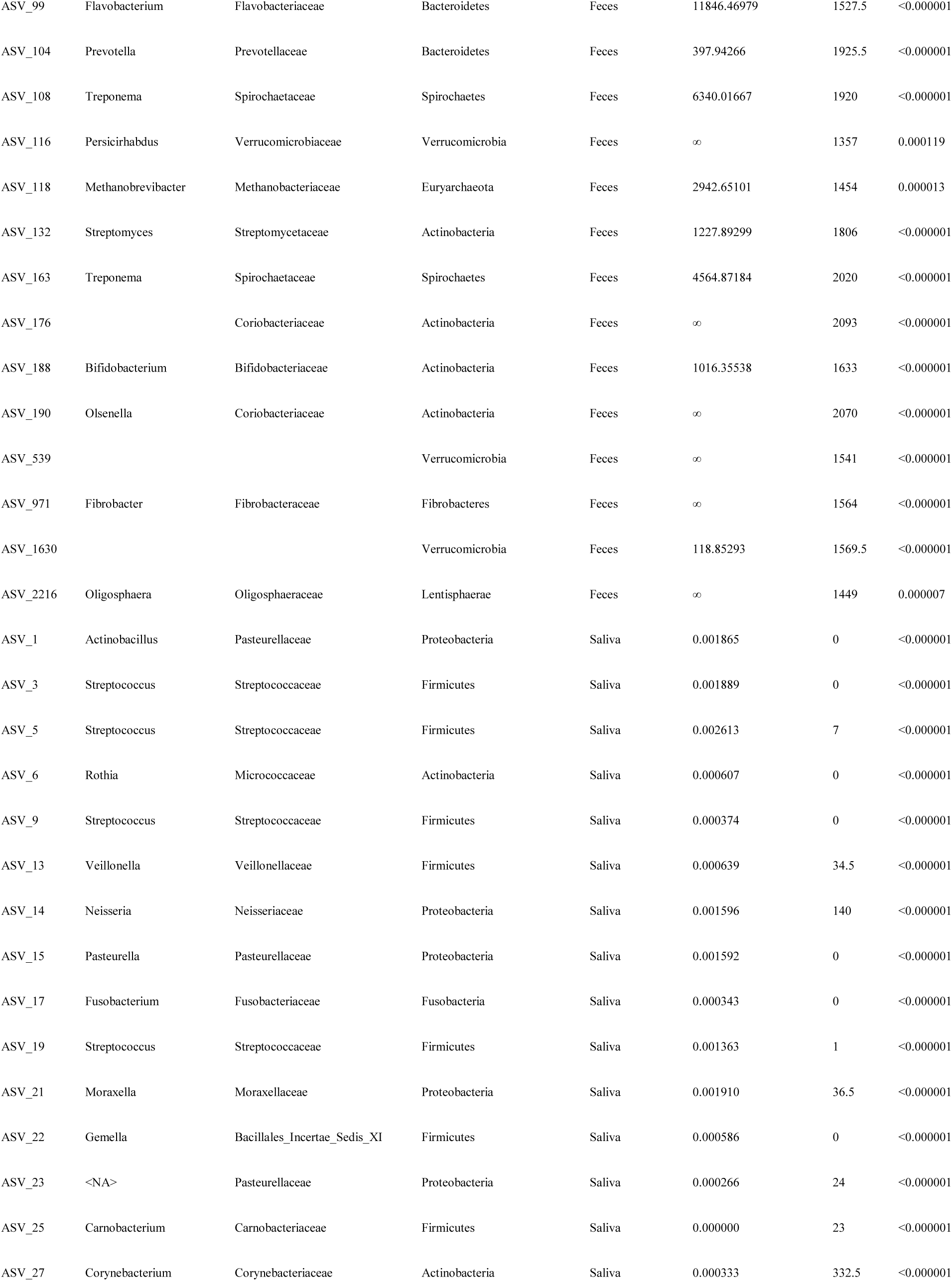
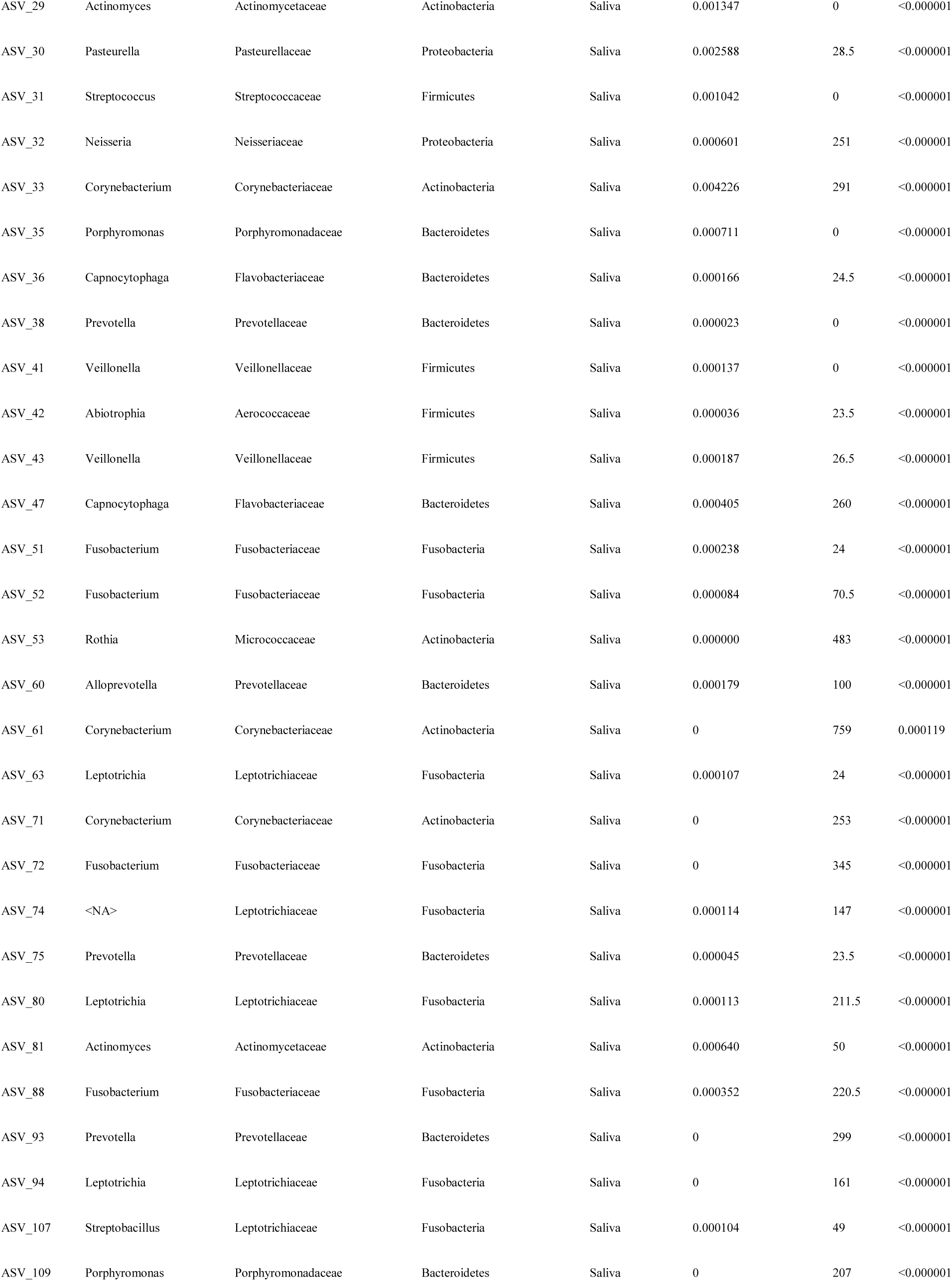
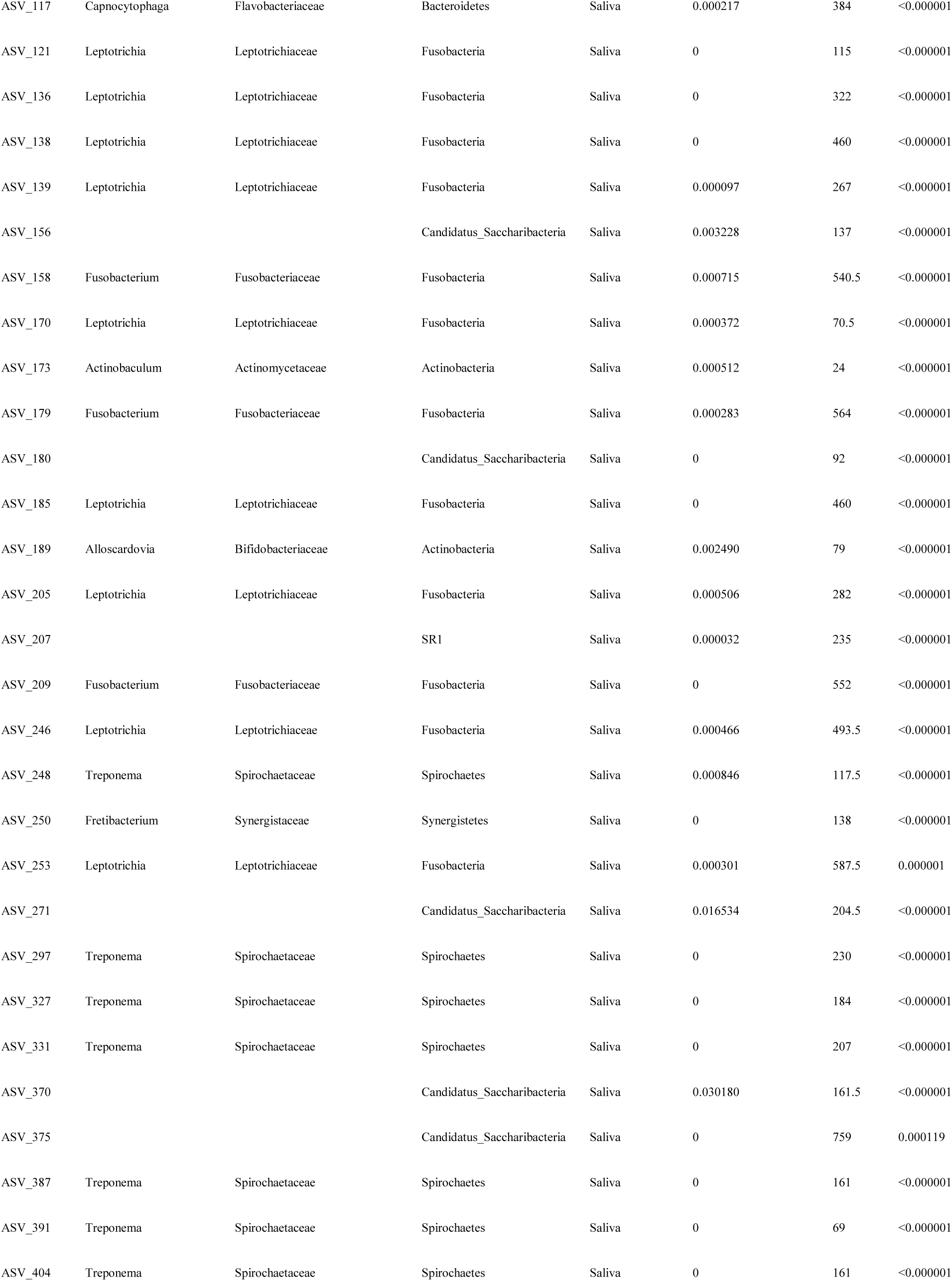
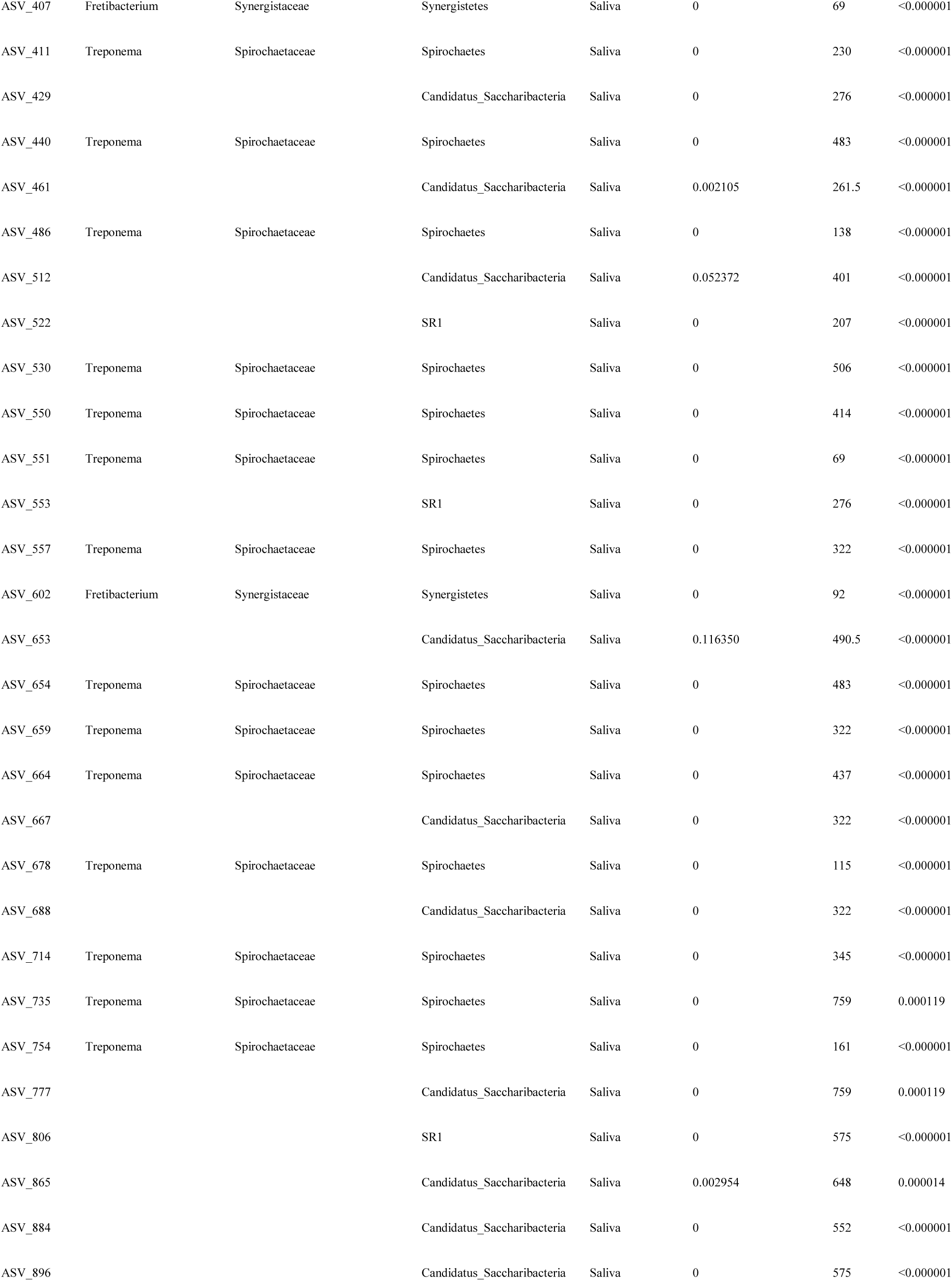
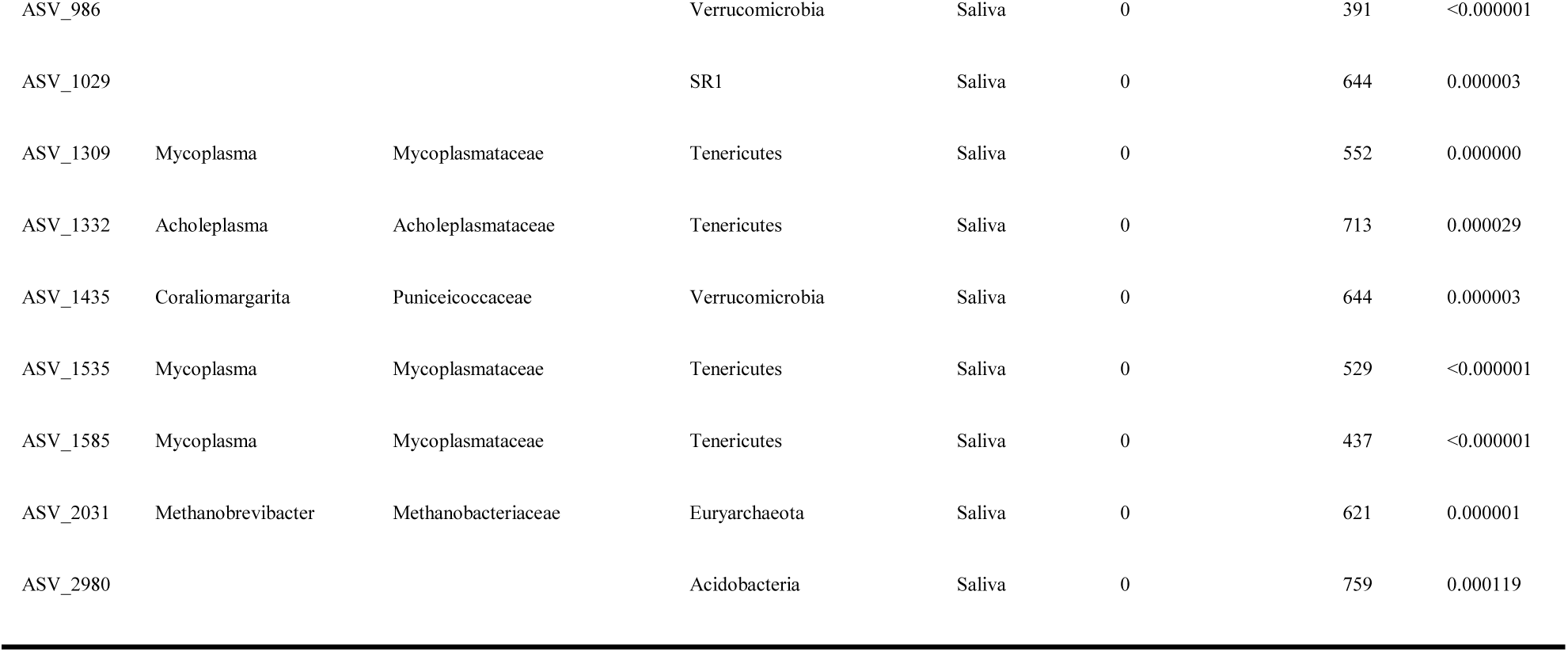
Taxonomic Differences in Composition of Saliva and Fecal Communities.

Thaumarchaeota (represented entirely by nitrogen-oxidizing families/ASVs)—were enriched in oral microbiomes. Each body site showed enrichment in similar numbers of families, with 14 families enriched in gut microbiomes relative to oral microbiomes and 15 families enriched in oral microbiomes relative to gut microbiomes. In contrast, oral microbiomes showed much higher enrichment for specific ASVs than gut microbiomes: only 33 ASVs were enriched in gut microbiomes relative to oral microbiomes, whereas 111 ASVs were enriched in oral microbiomes relative to gut microbiomes. Although too numerous to list in text, noteworthy gut-enriched ASVs matched to common Enterobacteriaceae, such as *Escherichia/Shigella*. *Prevotella* was also enriched in feces, as were *Pseudomonas* and *Treponema*. In contrast, oral microbiomes were heavily enriched in ASVs matching to *Streptococcus* (Phylum: Firmicutes), other strains of *Prevotella*, *Leptotrachia*, and their own distinct strains of *Treponema*.

We compared Shannon diversity, richness, evenness, and dispersion between gut and oral microbiomes (Supplementary Table S1). Only sampling locations including both fecal and saliva samples were compared (N=4). The oral microbiome showed higher Shannon diversity, richness, evenness at the ASV, family, and phylum level (Mann-Whitney-U tests: p<0.0001 for all tests) in comparison to the gut microbiome; whereas the gut microbiome showed higher dispersion at all three of these taxonomic levels (Permutation Test: p<0.0001 for all tests) compared to the oral microbiome.

Finally, we determined the taxonomic level at which variation for oral and gut microbiomes were most pronounced. Bray-Curtis distances across sampling locations showed lower AICc values at higher taxonomic levels for both gut (ASV: −42.78; Family: −67.19; Phylum: −108.16) and oral microbiomes (ASV: −81.86; Family: −119.14; Phylum: −146.67), suggesting that higher taxonomic levels explain more variation, relative to model (PERMANOVA) complexity for both saliva and fecal samples. However, total variation in Bray-Curtis dissimilarity between sites was explained by divergent taxonomic patterns in oral and gut microbiomes (Figure 4). In the case of oral microbiomes (Figure 5A), variation between collection sites was most pronounced at the level of ASVs (PERMANOVA: F= 2.57, R^2^= 0.357, Permutations=10000, p<0.0001), intermediate at the family level (PERMANOVA: F= 1.73, R^2^= 0.272, Permutations=10000, p=0.0194), and was not significantly variable at phylum level. In contrast, variation between collection sites for the gut microbiomes (Pearson Correlation: r= −0.999993, R^2^= 0.99999, p= 0.002; Figure 5B) was most pronounced at the phylum level (PERMANOVA: F= 3.72, R^2^= 0.266, Permutations=10000, p=0.005499), of intermediate importance at the family level (PERMANOVA: F=3.05, R^2^=0.229, Permutations=10000, p<0.0001), and least pronounced at the ASV level (PERMANOVA: F=1.98, R^2^=0.162, Permutations=10000, p<0.0001). No differences were seen in dispersion by site in either oral or gut microbiomes at any taxonomic level. Taken together, these results demonstrate that between site variation in oral microbiomes is dominated by smaller-scale variation of specific strains and families, whereas gut microbiomes tend to vary most consistently with one another at broad taxonomic groupings due to the concurrence of different constituent families and strains in co-localized samples.

**Figure 4.**
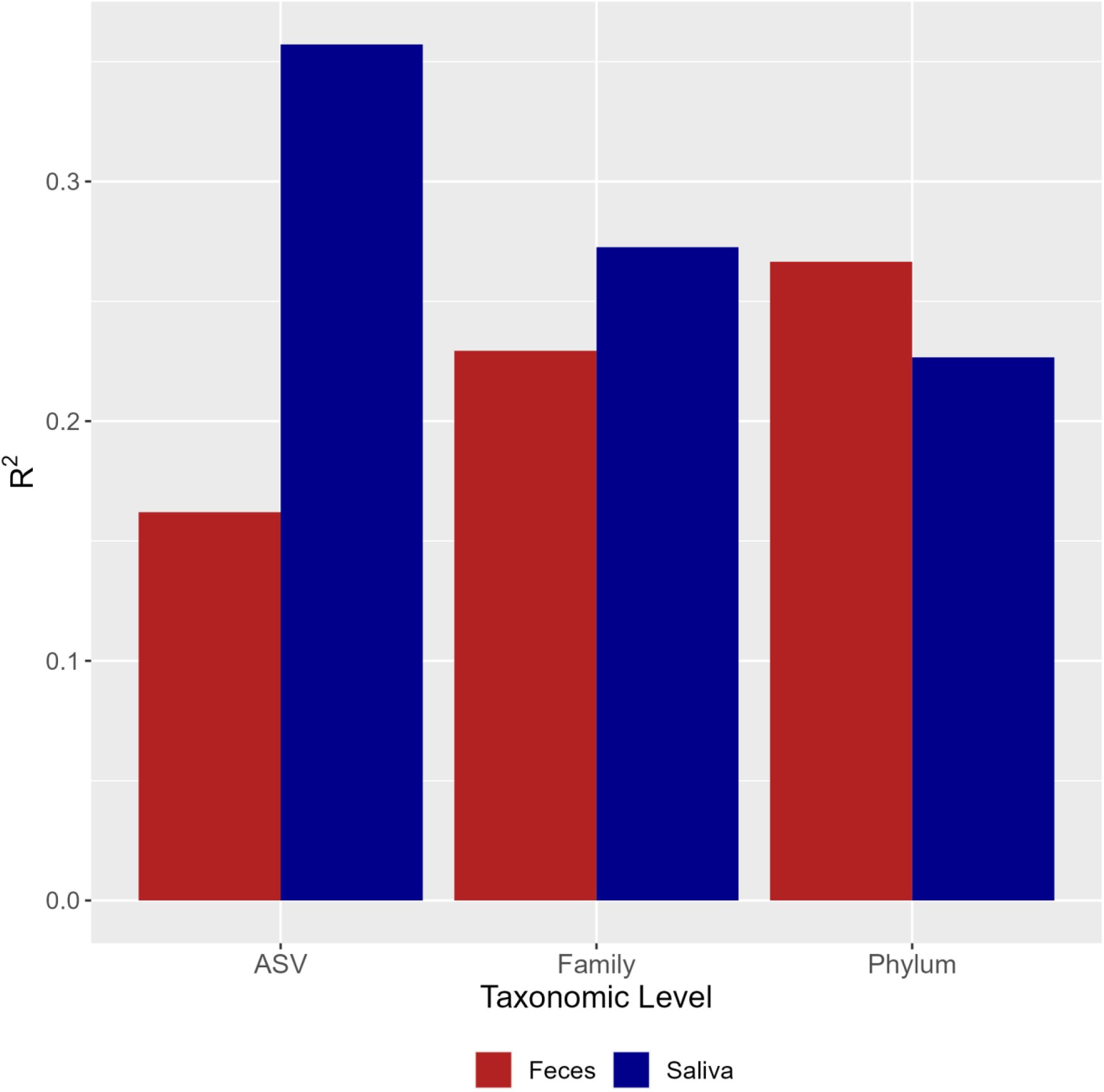
Between site variation explained by PERMANOVAs on Bray-Curtis dissimiliarties of gut and oral microbiomes at ascending taxonomic levels (left to right).

**Figure 5.**
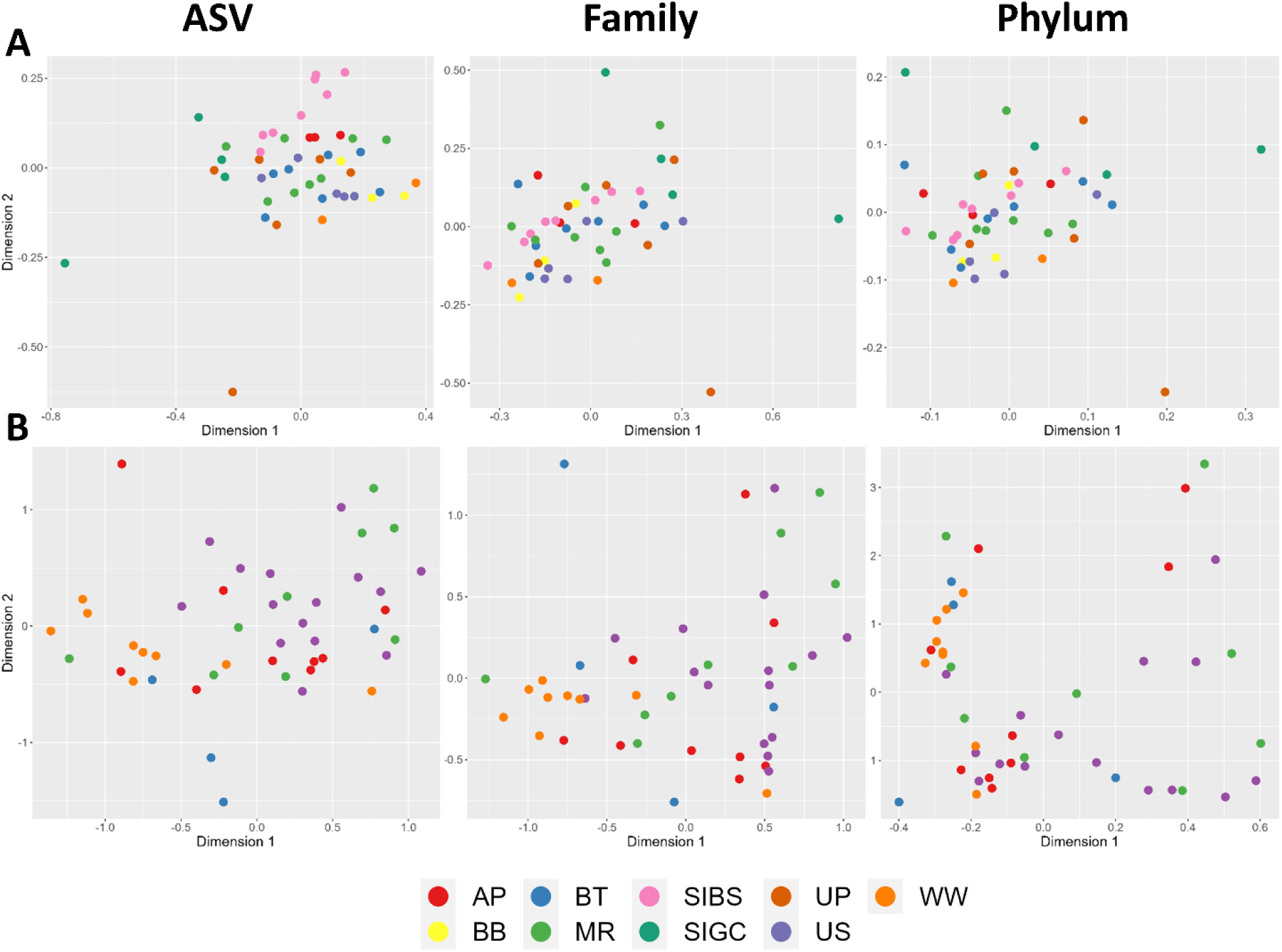
MDS of ASVs, families, and phyla (left to right) for saliva (A) and feces (B) colored by collection site. Stress for oral microbiomes are 16.73, 11.95, and 10.71 for ASVs, Families, and phyla, respectively. Stress for gut microbiomes are 20.11, 14.18, and 5.14 for ASVs, families, and phyla, respectively.

### Proteobacteria: Drivers of Taxonomic and Variation within Body Sites

We assessed variation in body sites beginning at the levels suggested to be of greatest importance in explaining between site variation across samples. In the case of saliva samples, this was the ASV level. In the case of the gut microbiome, we began at a broad phyla level and then narrowed this down to the particular families and functional ASVs within these identified phyla.

We first used PCA to assess the phyla that drove variation in relative bacterial assemblies within each body site (Figure 6A-B). Three ASVs played a predominant role in explaining variation in oral microbiome: ASV-1 (Genus: *Actinobacillus*; Phylum: Proteobacteria; Variance Explained: 20.0%), ASV-5 (Genus: *Streptococcus*; Phylum: Firmicutes; Variance Explained: 31.3%), and ASV-61 (Genus: *Corynebacterium*; Phylum: Actinobacteria; Variance Explained: 18.7%). Clear groupings were not self-evident in oral microbiomes even at the ASV level, although a group of four samples did appear to show some distinction based on the relative abundance of ASV-5 (Figure 6A). In the gut microbiome, variation in assembly structure was primarily explained by an antagonistic relationship in the relative abundances of Proteobacteria and Firmicutes. Bacteroidetes was a minor driver of assembly structure in the gut microbiome. Proteobacteria explained the largest proportion of variation (46.3%) in the gut microbiome, followed by Firmicutes (42.8%) and then Bacteroidetes (8.91%). Actinobacteria explained a minor portion of variation in the gut microbiome (1.5%), and no other phyla explained more than 1% of variation.

**Figure 6.**
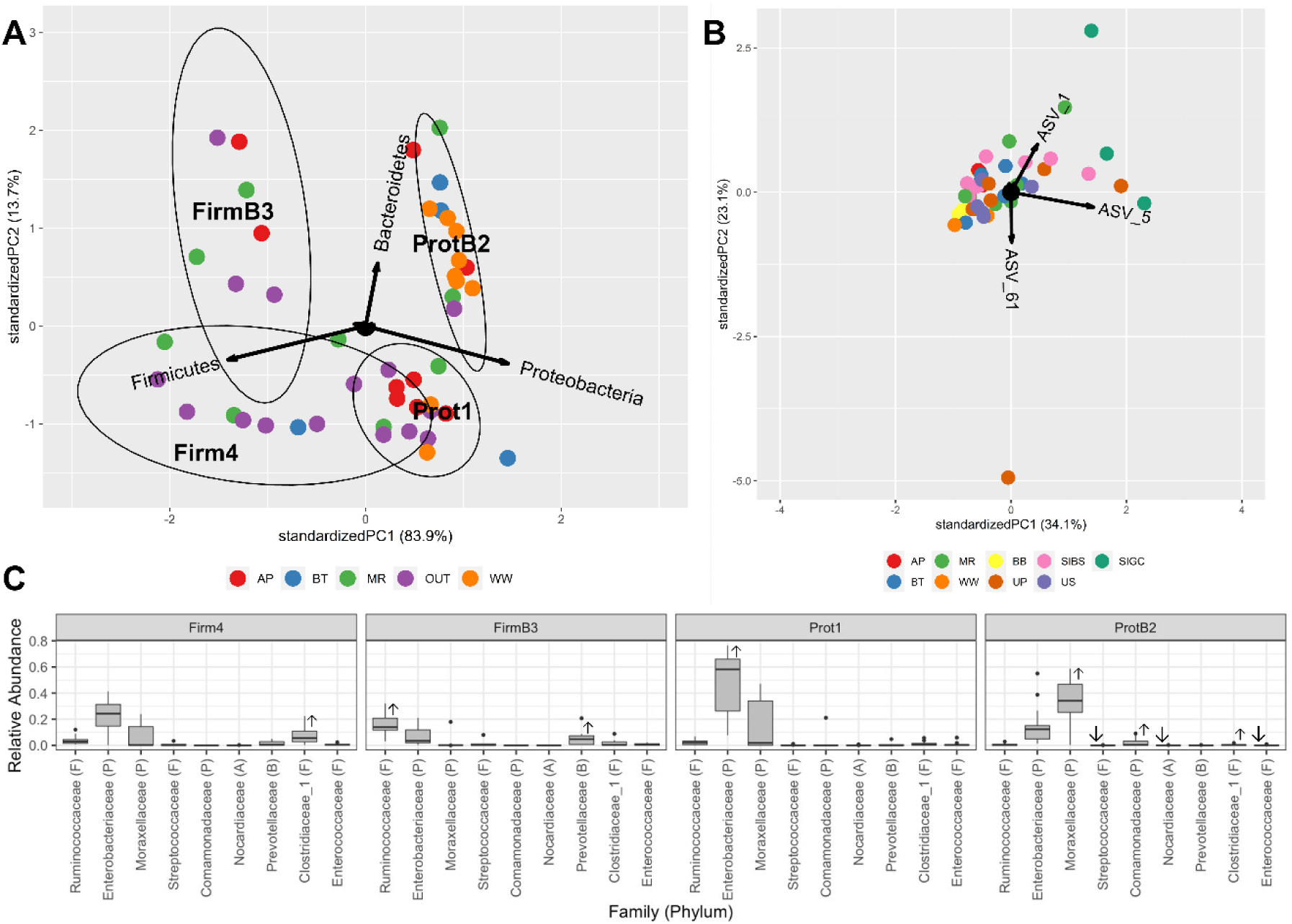
PCA biplots of oral ASV and (A) and (B) gut phyla microbiome compositions. Correlations of major taxa with the principal components are shown as black arrows with longer arrows corresponding to stronger correlations. Samples are color-coded by site and variance explained by principal components is shown on their respective axis. Groups are assigned based on the sign of the principal components. Ellipses show 95% confidence intervals for clusters based on an assumption of a multivariate t-distribution. Families with significant differences between major clusters in the gut PCA are shown as box plots (C). Families that are significantly enriched (↑) or depleted (↓) in each cluster are denoted with an arrow. ASV_1 corresponds to *Actinobacillus* (Proteobacteria); ASV_5 corresponds to *Streptococcus* (Firmicutes); and ASV_61 corresponds to *Corynebacterium* (Actinobacteria).

Ultimately, the gut microbiome appeared to show 3-4 distinct groupings of samples in its 2D PCA (Figure 6B), largely based on the sign of its standardized principal components. A clear division was seen between high Proteobacteria (ProtB2) and high Firmicutes (FirmB3) samples among samples higher in Bacteroides (PC2>0). These two groupings were also clearly distinct from samples with a lower relative abundance of Bacteroidetes (PC2<0). However, groupings were less distinct among samples with lower relative abundances of Bacteroidetes, despite strong stratification—and compositional differences—based on the relative abundances of Proteobacteria and Firmicutes.

After finding groupings of microbiome communities in gut microbiomes based on phyla, we used Mann-Whitney U Tests to identify the constituent families driving differences between these groups. A Dunn-Šidák corrected of ɑ = 0.00107 of was used for these tests based on the presence of 48 families (Figure 6C). All groups showed variation in abundances at the family level (Figure 6C). Samples dominated by Proteobacteria, but were low in Firmicutes and Bacteroidetes (Prot1), were enriched in the Proteobacteria family Enterobacteriaceae (p=0.00018) relative to other groups. Samples characterized by higher abundances of Proteobacteria and Bacteroidetes (ProtB2) were enriched in the Proteobacteria families Comamonadaceae (p= 0.00097), and Moraxellaceae (p=0.00012) but depleted in the Firmicutes families Enterococcaceae (p=0.00087) and Streptococcaceae (p=0.00012), as well as the Actinobacteria family Nocardiaceae (p=0.00106). Families high in Firmicutes and Bacteroidetes (FirmB3), but low in Proteobacteria, were enriched in the Firmicutes family Ruminococcaceae (p=0.00013) and the Bacteroidetes family Prevotellaceae (p=0.00085). Samples high in Firmicutes but low in Proteobacteria and Bacteroidetes (Firm4) were enriched in Clostridia-1 (p=0.00034).

As gut microbiomes were structured based on Proteobacteria, Firmicutes and Bacteroidetes, we assessed the functional implications of the Proteobacteria:Firmicutes (P:F) ratio in comparison to the traditionally-defined Firmicutes:Bacteroidetes (F:B) ratio by running the ASV data through PiCrust2 with Aldex. We predicted 22 differentially expressed metabolic pathways based on the Proteobacteria to Firmicutes ratio (Table 3). In total, 18 of these pathways were enriched in samples with more Proteobacteria than Firmicutes and primarily related to lipid and fatty acid biosynthesis and degradation of potentially harmful aromatic compounds, such as catechol and salicylate. In contrast, only four pathways were enriched in high Firmicutes and low Proteobacteria samples with two of these relating to vitamin B12 production. These differences corresponded to enrichment in the Proteobacteria families Moraxellaceae (p=0.0009) and the Actinobacteria families Nocardiaceae (p=0.0009) and Streptomycetaceae (p=0.0008) in sample dominated by Proteobacteria, and the Actinobacteria family Coriobacteriaceae (p=0.00003), the Bacteroidetes family Prevotellaceae (p=0.0003), and four families of Firmicutes— Erysipelotrichaceae (p=0.00005), Lachnospiraceae (p=0.0001), Lactobacillaceae (p=0.00003), Ruminococcaceae (p=0.0003)—in samples in which Firmicutes dominated Proteobacteria. We found 10 metabolic pathways that were differentially expressed between Firmicutes and Bacteroidetes dominated samples in the gut (Table 4). These were equally split in enrichment among higher Firmicutes and higher Bacteroidetes samples. Three of the five enriched pathways in samples with higher Firmicutes were related to pyridine or DNA production, whereas four of the five pathways enriched in samples with higher Bacteroidetes were involved in the degradation of simple organic compounds (fructose, pyruvate, and glycerol). These differences corresponded to enrichment of three families of Firmicutes within samples in which Firmicutes dominated Bacteroidetes: Ruminococcaceae (p=0.00026), Planococcaceae (p=0.00025), and Lachnospiraceae (p<0.00001) and enrichment of two Bacteroidetes families Sphingobacteriaceae (p=0.00003) and Flavobacteriaceae (p<0.00001) in samples in which Bacteroidetes-rich dominated Firmicutes. No differences were found between Bacteroidetes and Firmicutes dominated saliva samples, although we had only one saliva sample with more Bacteroidetes than Firmicutes.

**Table 2.**
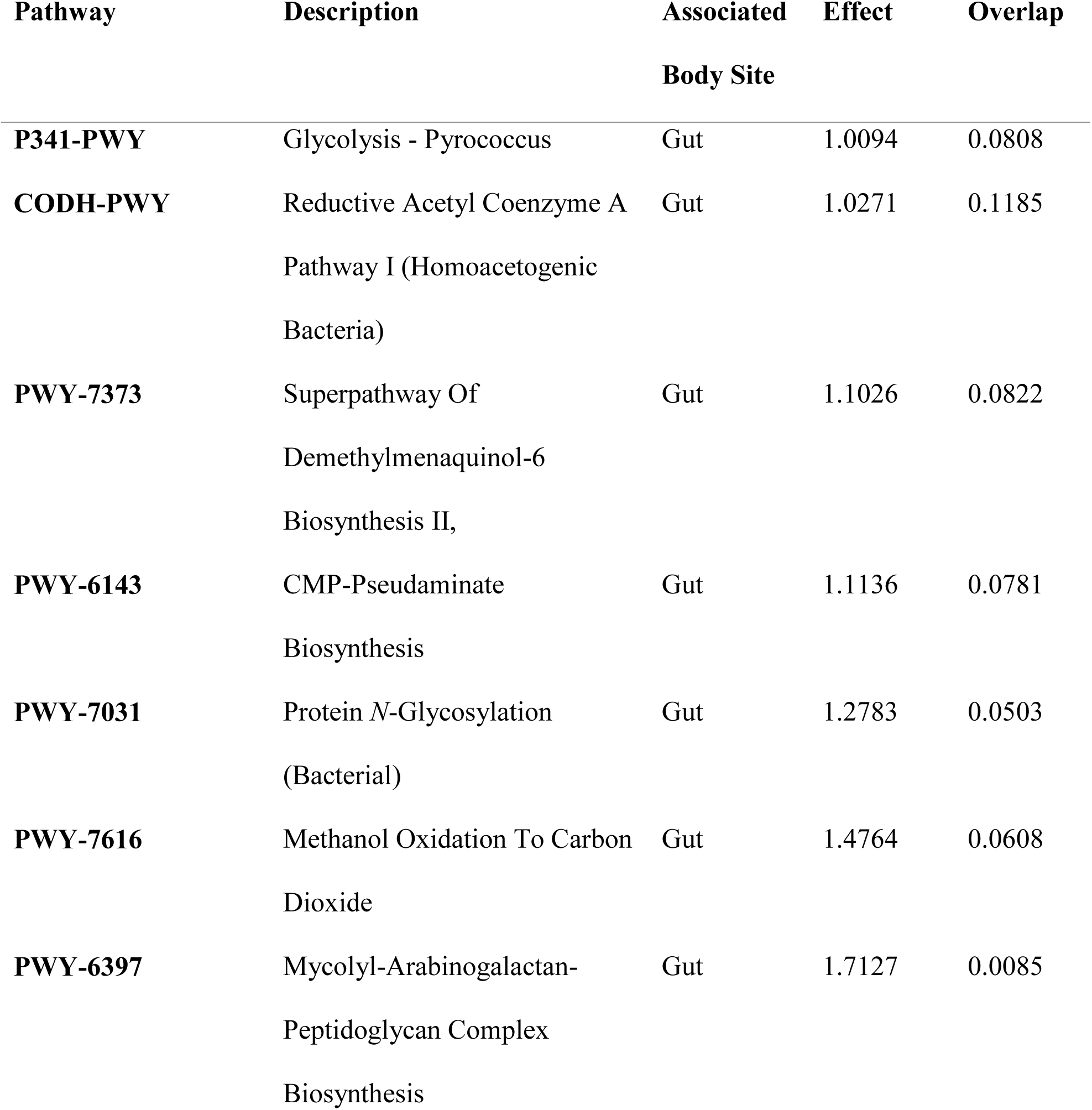
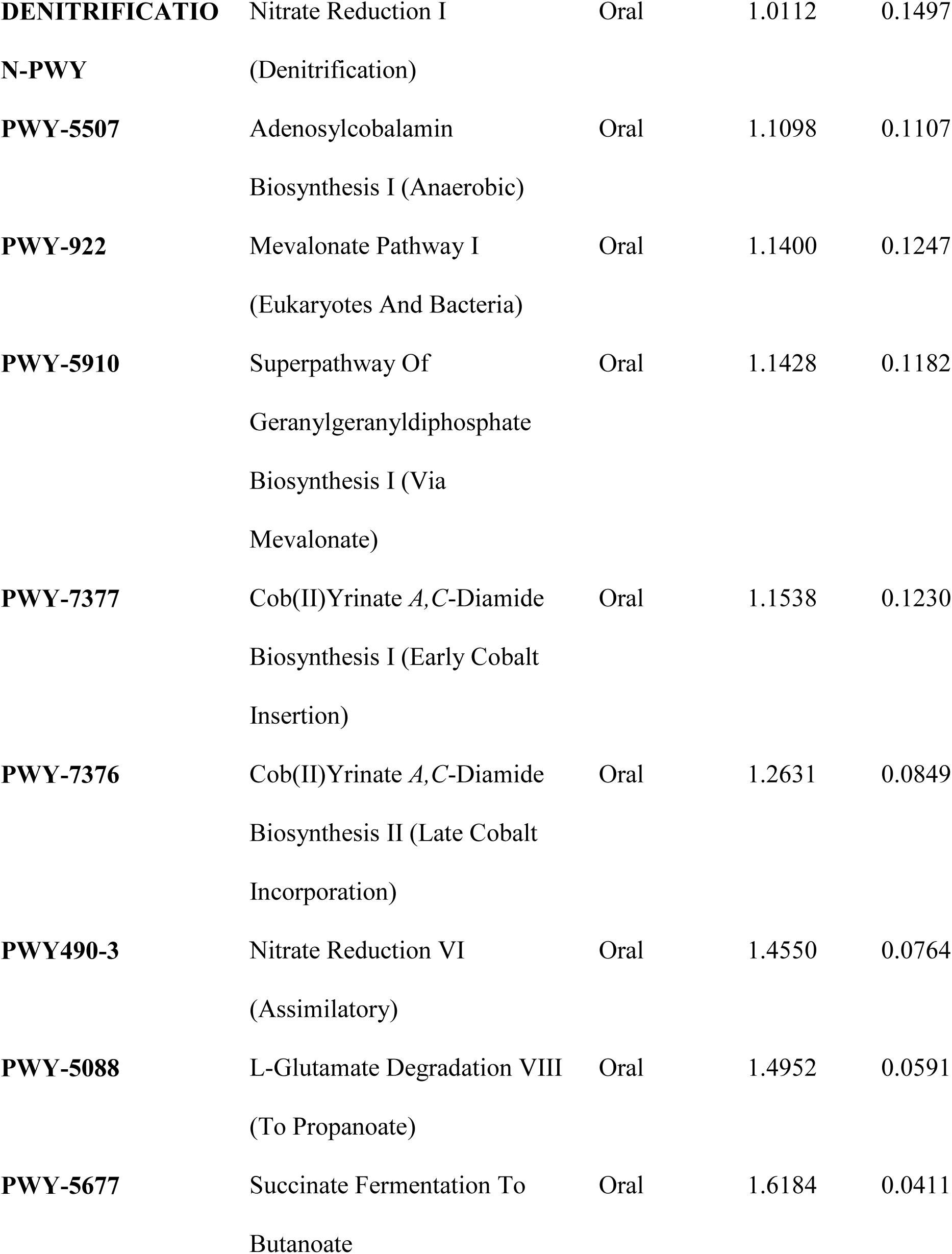
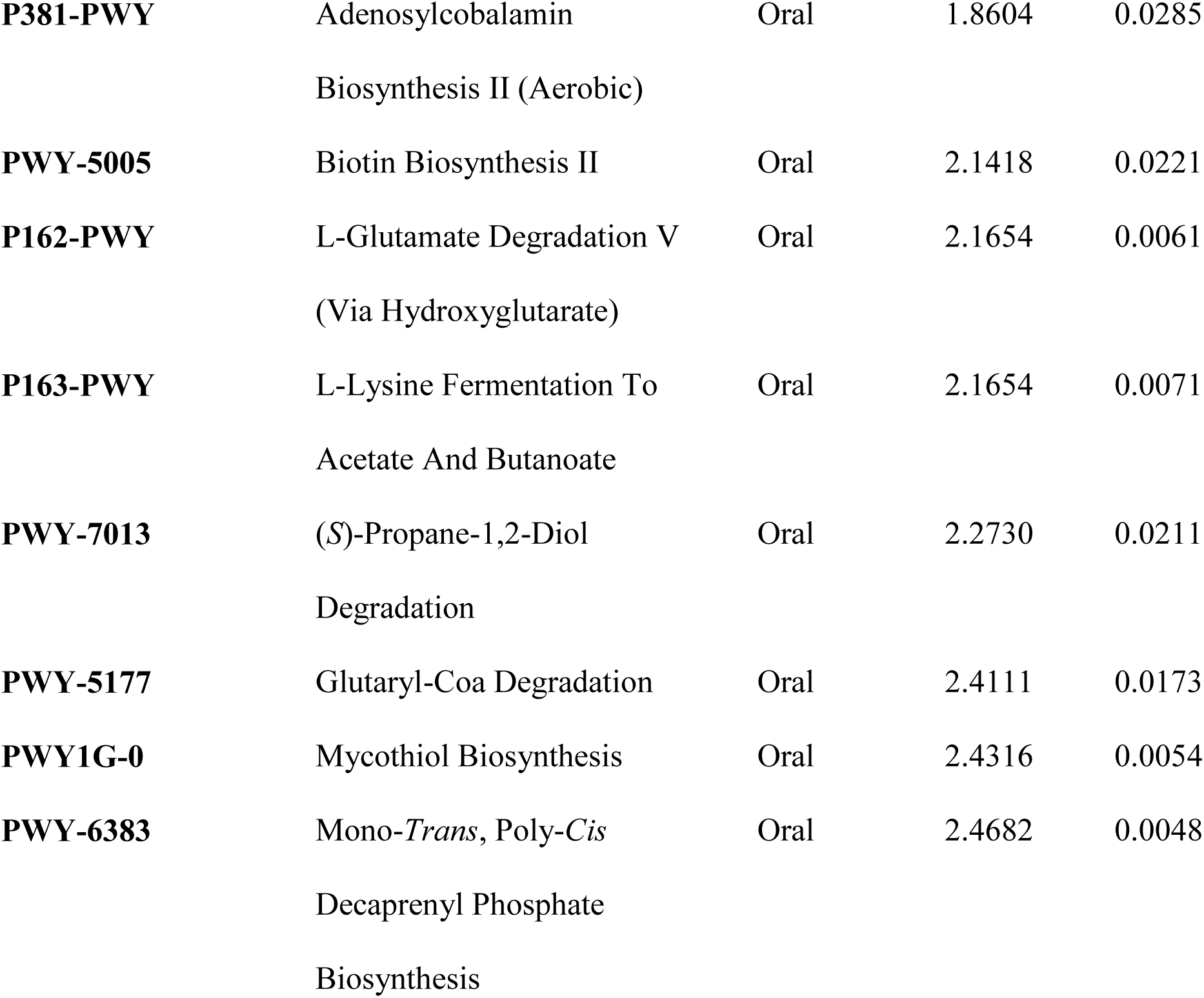
Predicted MetaCyC pathways with an effect size of > 1.0 as estimated by Aldex2 - comparison for gut or oral microbiomes.

**Table 3.** Predicted MetaCyC pathways with an effect size of > 1.0 as estimated by Aldex2 – comparison for Proteobacteria- or Firmicutes-rich gut microbiomes.

| Pathway | Description | Dominant<br>Taxa | Effect | Overlap |
| --- | --- | --- | --- | --- |
| **BIOTIN-**<br>**BIOSYNTHESIS-**<br>**PWY** | biotin biosynthesis I | Proteobacteria | 1.0795 | 0.109375 |
| **CATECHOL-**<br>**ORTHO-**<br>**CLEAVAGE-PWY** | catechol degradation to β-<br>ketoadipate | Proteobacteria | 1.0437 | 0.131113 |
| **FASYN-INITIAL-**<br>**PWY** | superpathway of fatty acid<br>biosynthesis initiation (E. coli) | Proteobacteria | 1.1478 | 0.066667 |
| **KDO-**<br>**NAGLIPASYN-**<br>**PWY** | superpathway of (Kdo)2-lipid<br>A biosynthesis | Proteobacteria | 1.1799 | 0.108221 |
| **NAGLIPASYN-**<br>**PWY** | superpathway of (Kdo)2-lipid<br>A biosynthesis | Proteobacteria | 1.2300 | 0.082206 |
| **PPGPPMET-PWY** | ppGpp biosynthesis | Proteobacteria | 1.3121 | 0.094693 |
| **PROTOCATECHU**<br>**ATE-ORTHO-**<br>**CLEAVAGE-PWY** | protocatechuate degradation II<br>(ortho-cleavage pathway) | Proteobacteria | 1.0885 | 0.12487 |
| **PWY-1269** | CMP-3-deoxy-D-manno-<br>octulosonate biosynthesis I | Proteobacteria | 1.2294 | 0.075 |
| **PWY-5417** | catechol degradation III (ortho-cleavage pathway) | Proteobacteria | 1.1641 | 0.115505 |
| **PWY-5431** | aromatic compounds degradation via β-ketoadipate | Proteobacteria | 1.1402 | 0.114583 |
| **PWY-6182** | superpathway of salicylate degradation | Proteobacteria | 1.0877 | 0.127083 |
| **PWY-6185** | 4-methylcatechol degradation (ortho cleavage) | Proteobacteria | 1.0642 | 0.1375 |
| **PWY-6282** | palmitoleate biosynthesis I (from (5Z)-dodec-5-enoate) | Proteobacteria | 1.1799 | 0.084287 |
| **PWY-6519** | 8-amino-7-oxononanoate biosynthesis I | Proteobacteria | 1.1752 | 0.096774 |
| **PWY-7664** | oleate biosynthesis IV (anaerobic) | Proteobacteria | 1.0967 | 0.097917 |
| **PWY0-42** | methylcitrate cycle I | Proteobacteria | 1.1826 | 0.104058 |
| **PWY0-845** | superpathway of pyridoxal 5'-phosphate biosynthesis and salvage | Proteobacteria | 1.3592 | 0.082292 |
| **UBISYN-PWY** | superpathway of ubiquinol-8 biosynthesis (prokaryotic) | Proteobacteria | 1.1722 | 0.095833 |
| **CENTFERM-PWY** | pyruvate fermentation to butanoate | Firmicutes | -1.0103 | 0.126951 |
| **PWY-5088** | L-glutamate degradation VIII<br>(to propanoate) | Firmicutes | 1.1144 | 0.135417 |
| **PWY-5507** | adenosylcobalamin<br>biosynthesis I (early cobalt<br>insertion) | Firmicutes | 1.0548 | 0.11875 |
| **PWY-7377** | cob(II)yrinate a,c-diamide<br>biosynthesis I (early cobalt<br>insertion) | Firmicutes | -1.1699 | 0.120833 |

**Table 4.**
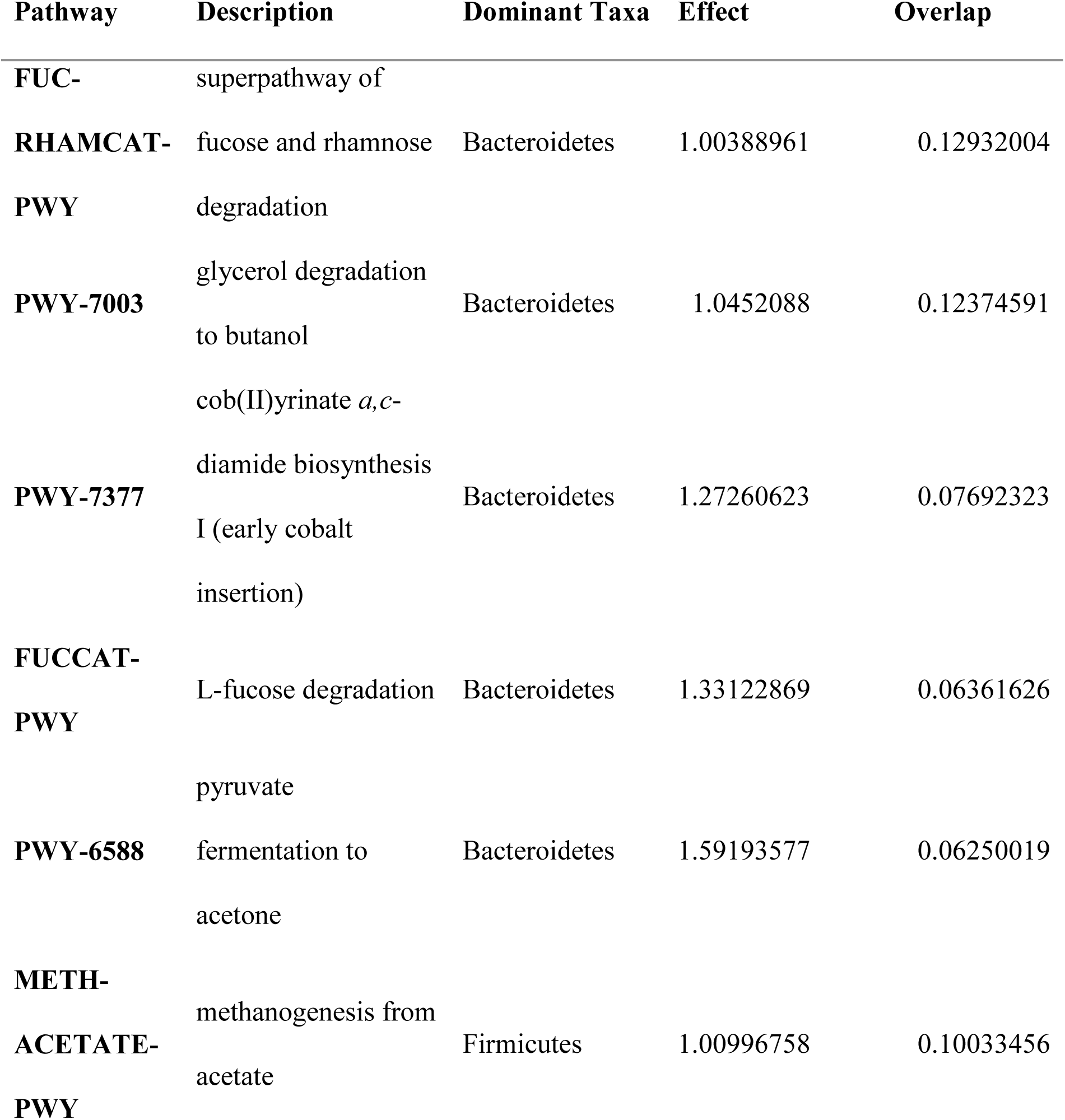
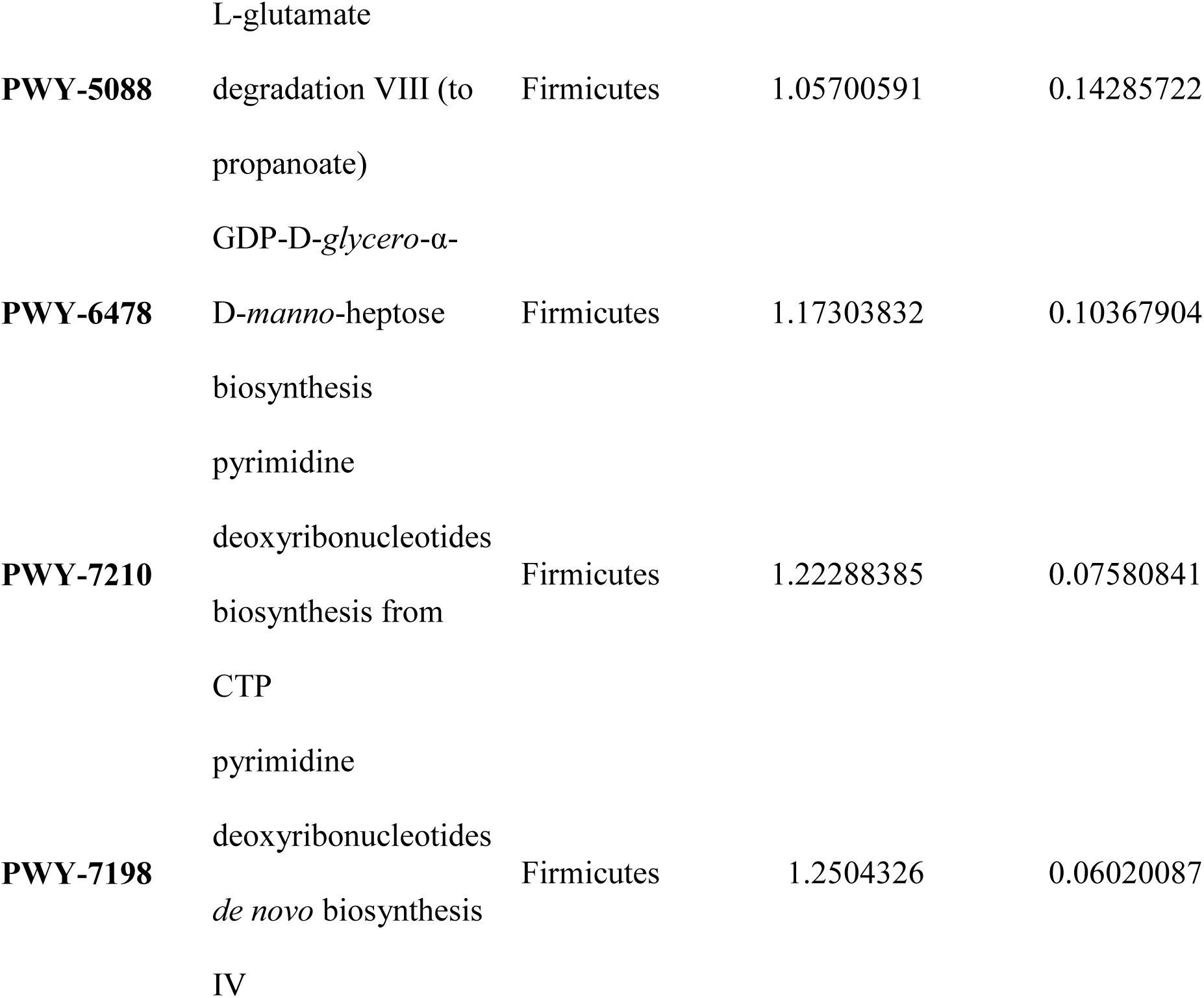
Predicted MetaCyC pathways with an effect size of > 1.0 as estimated by Aldex2 – comparison for Firmicutes- or Bacteroidetes-rich gut microbiomes.

## DISCUSSION

Here we analyzed the oral and gut microbiome of wild long-tailed macaques in Singapore using metabarcoding of the V4 hypervariable region of the 16S gene with the goal of characterizing the bacterial and archaeal communities of an urban non-human primate. We find that the oral and gut bacterial microbiomes were both dominated by Proteobacteria, Firmicutes, and Bacteriodetes. Archaea were rare but consisted overwhelmingly of methanogens at both body sites. Methanogens were in fact the only archaea found in feces, but numerous clades of ammonia-oxidizing (nitrifying) archaea were found in saliva. Saliva microbiomes had higher α-diversity than gut microbiomes and were more even and richer at all taxonomic levels. In contrast, gut microbiomes showed greater dispersion in β-dissimilarity across samples at all levels. Structure within oral and gut microbiomes occurred in contrasting taxonomic patterns, with more pronounced between-site variation at lower taxonomic units in the oral microbiomes, and more pronounced between site variation at higher taxonomic units among fecal samples. Distinct groupings of oral microbiomes were not seen at any taxonomic level, but gut microbiomes did break into 3-4 distinct groups at the phylum level. Groupings within gut microbiomes were explained by relative abundances of Proteobacteria, Firmicutes, and Bacteroidetes. Proteobacteria and Firmicutes accounted for similar proportions of taxonomic variation between samples, but Proteobacteria accounted for much more inferred functional differentiation: microbiomes high in Proteobacteria were enriched in >4X more pathways than microbiomes high in Firmicutes and the P:F ratio explained almost twice as many functional differences as the F:B ratio.

Our analyses are the first microbiome annotations of saliva and feces for urban long-tailed macaques and complement existing annotations from captive and wild non-urban macaques in Thailand (Grant et al., 2019; Sawaswong et al., 2021). We report some similarities to these previous annotations: saliva and gut microbiomes were clearly distinct; the same phyla, families and ASVs are generally present at respective body sites; and gut microbiomes show decidedly higher levels of ß-diversity differentiation from one another. However, the differences between annotations are more striking, Singapore’s macaques show: decidedly higher Shannon diversity in oral microbiomes than gut microbiomes at lower taxonomic levels; a dominant role for Firmicutes in the oral microbiome, but a dominant role for Proteobacteria in the gut; and much greater variation in assembly structure within both body sites. In contrast, reports from Thailand (Sawaswong et al., 2021) indicated: higher diversity (α and ß) in the saliva relative to feces in captive macaques only; a dominant role for Firmicutes in feces and Proteobacteria in saliva; and fairly consistent patterns of assembly structure for both oral and oral and gut microbiomes— whether wild or captive. Overall, these comparisons demonstrate that within species differences in microbiome can be immense between sampling populations, habitats, and localities.

We also provide the first characterization of archaea in long-tailed macaques. These archaeal communities generally resemble those of humans with a dominant role for methanogens in the oral cavity and gut, and a limited presence of nitrifying Thaumarchaeota in the oral cavity as well (Weiland-Bräuer, 2023). Indeed, methanogens are the only known archaea in the human gut microbiome (Nishida & Ochman, 2019), and were the only archaea found in the gut microbiomes of Singapore’s macaques. In contrast, both Thaumarchaeota and methanogens have been reported in feces of great apes (Nishida & Ochman, 2019). Archaeal communities have been reported to correlate primarily with host phylogenies in vertebrates and to a lesser extend with host diet (Youngblut et al., 2021). The samples that we studied showed sufficient number of families and ASVs in both the gut and oral cavities to be variable with regard to diet. Changes in archaeal communities based on diet may be of interest due to the role of these symbionts in both carbon cycling and environmental modelling within the gut (Weiland-Bräuer, 2023). Notably, we see a predicted enrichment of methanogenosis pathways in Firmicutes-rich samples of the gut, suggesting that dietary changes may influence both archaeal composition and production of methane. If high compositions of Firmicutes do in fact represent a ‘westernization’ (Malesza et al., 2021; Ross et al., 2024) of macaque diet, then urban development or other anthropogenic factors may drive greenhouse gas production by macaques. However, the specific drivers of gut microbiome assembly structure in macaques remain unclear, and the application of such human-centric interpretations has been called into question (Amato et al., 2015). Our present study provides only a tentative first look at archaeal communities of long-tailed macaques in light of only low level amplification of archaea in our samples and the high likelihood that existing databases are lacking in archaeal reference reads (Manara et al., 2019). Our findings nonetheless suggest that Singapore’s macaques possess oral and gut archaeal communities that resemble those of humans in both their taxonomy and functional roles.

Bacterial communities were highly variable within the gut and showed groupings that were most pronounced at the phylum level. These groups also displayed some correlation to collection site and were also correlated with functional differences. These emergent patterns of variation at higher taxonomic levels suggest the presence of broadly shared and cohesive ecological and functional pathways within these taxa (Philippot et al., 2010), in the context of highly stochastic colonization or highly variable localized niche space within the guts of individual macaques. It is plausible then that variability in gut microbiomes may provide a source of ecological plasticity to Singapore’s macaques. Indeed, the clear variability of gut microbiome composition in Singapore’s macaques relative to those in Thailand (Grant et al., 2019; Sawaswong et al., 2021) may reflect more highly variable food sources and environments within Singapore’s urban landscape (Sha & Hanya, 2013), consistent with the finding that human exposure increased variability of macaque microbiomes in Thailand as well (Grant et al., 2019). Taxonomic and functional variation do not, however, automatically imply variation in other aspects of host phenotype and fitness (Brüssow, 2020). We have no measurements on macaque health, and lack of knowledge of conserved impacts of microbiota by hosts in NHPs preclude additional inferences into the ultimate adaptive potential of microbial variation in Singapore’s macaques.

The gut microbiomes of Singapore’s macaques were in fact quite distinct from those observed in humans and most other NHPs (Clayton et al., 2018). Though similar phyla and taxa were present, the dominance of Proteobacteria over Firmicutes and Bacteroidetes is unusual. The presence of Proteobacteria provide an additional dimension of taxonomic and functional variation within Singapore’s macaques. Microbiomes low in Proteobacteria follow classical patterns associated with the F:B ratio in primates: high proportions of Bacteroidetes reflect enrichment of Prevotellaceae and resemble assemblies from humans consuming non-western diets rich in plant matter; and high proportions of Firmicutes reflect enrichment in Clostridiaceae_1 (primarily *Clostridium*) and resemble assemblies from humans consuming ‘westernized’ diets rich in fat (Malesza et al., 2021; Ross et al., 2024). The P:F ratio, however, predicts far more variation in both assembly structure and inferred function of Singapore’s macaques. The biological significance of this variation may have far-reaching implications for the role of Proteobacteria as sources of adaptability or as indicators of health.

Proteobacteria may represent novel sources of host plasticity (Bradley & Pollard, 2017) or threats to host health (Rizzatti et al., 2017). Gut microbiomes high in Proteobacteria were enriched in either Moraxellaceae or Enterobacteriaceae, both of which generally reflect dysbiosis in humans (Rizzatti et al., 2017). High levels of Proteobacteria have also been associated with *Plasomodium* infection in rhesus macaques (Farinella et al., 2023) and humans (Winaris et al., 2023). While we can confirm that *Plasmodium* infection is effectively universal in Singapore’s long-tailed macaques (Wilcox *et al*., 2018), dysbiosis of the gut is often characterized by loss of function (Kriss et al., 2018; VanEvery et al., 2023). In contrast, our detection of the heavily skewed distribution towards gain of function in Proteobacteria-dominated microbiomes suggests that they may play ecologically and metabolically important roles within Singapore’s long-tailed macaques. Whether these functions provide any use to their hosts remains totally undetermined: more diverse profiles have been associated with healthier phenotypes in the human gut (VanEvery et al., 2023), but there are exceptions (Brüssow, 2020). High levels of Enterobacteriaceae are suggested to provide many unique and individualized functions in the human gut (Bradley & Pollard, 2017). Our findings of substantial enrichment in degradation and biosynthesis pathways associated with Proteobacteria are consistent with this finding. Our findings therefore contribute to existing literature, suggesting potentially important ecological and functional significance to high levels of Proteobacteria in an NHP.

Ultimately our findings demonstrate additional sources of variation in the microbiomes of long-tailed macaques relative to their conspecifics in Thailand (Grant et al., 2019; Sawaswong et al., 2021) and humans (Ross et al., 2024). Major questions remain as to the capacity for cross-species inferences in microbiome research on primates. Our findings show that interspecific variation within fecal microbiome assemblages of long-tailed macaque can be immense with considerable variation in even the dominant phyla that are present. Such variation may provide for important ecological plasticity or conversely indicate new sources of pathology. It also complicates both inter- and intraspecific inferences into the implications of variation in microbiome structure. The capacity for Proteobacteria and the P:F ratio to predict more variation than the F:B ratio in our system raises questions as to the roles and implications of all three bacterial phyla. Although Firmicutes still certainly provide a capacity for lipid degradation under anaerobic conditions, they may no longer provide biologically informative information on lipid consumption if they are competing with Proteobacteria capable of assuming similar functions. As we show, such variation may also only be apparent if examined across many individuals, foraging across diverse environmental and geographic space. It may likewise be missed altogether or be more apparent at specific taxonomic levels of constituent taxa. Broad characterizations of primate microbiomes across environments and taxonomic space are therefore still required in order to build this more complete picture of patterns of variation within species. Long-tailed macaques provide an excellent system for the study of microbiome ecology across space, scales, and anthropogenic landscapes.

## ACKNOWLEDGMENTS

This work was funded by the National Science Foundation (NSF) Division of Behavioral and Cognitive Sciences (BSC 0629787), the NSF IGERT GLOBES program (#0504495), the National Science Foundation East Asian Pacific Summer Institute Singapore program, and the following groups within the University of Notre Dame: the Institute for Scholarship in the Liberal Arts, the Center for Aquatic Conservation, and the College of Science. This research was supported in part by the Notre Dame Center for Research Computing through use of the high-performance computing cluster. The Notre Dame Genomics & Bioinformatics Core Facility assisted through sequencing consultation and library preparation. We specifically acknowledge the contributions of Jaqueline Lopez and Melissa Stephens in the genomics core facility for their advice and helpful suggestions regarding sequencing. We would like to acknowledge Dr. Amy Klegarth for assistance in data collection and sample processing.

## COMPETING INTERESTS

The authors declare that they have no competing interests.

## DATA AVAILABILITY

All sequencing data are available on Sequence Read Archive of the National Center for Biotechnology Information of the National Institutes of Health National Library of Medicine (project Accession PRJNA776160).

## Notes

https://www.ncbi.nlm.nih.gov/bioproject/?term=(PRJNA776160)%20AND%20bioproject_sra[filter]%20NOT%20bioproject_gap[filter]

### Competing Interest Statement

The authors have declared no competing interest.

