## Supplementary figures and images for "Characterizing Bacterial and Archaeal Microbiomes of Singapore’s Urban Long-Tailed Macaques"

### Supplementary Figure 1

#### Saliva Samples

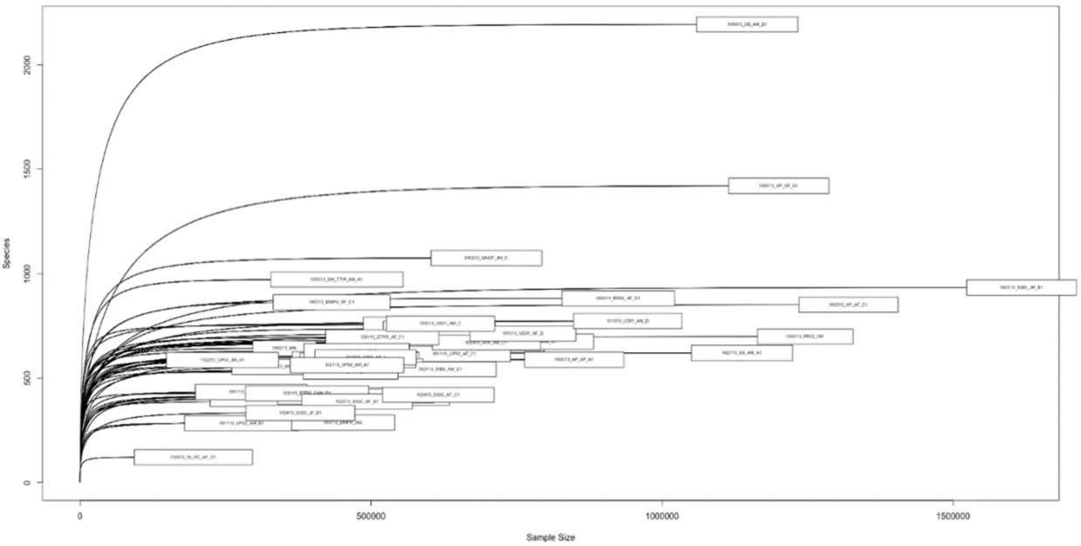


#### Fecal Samples

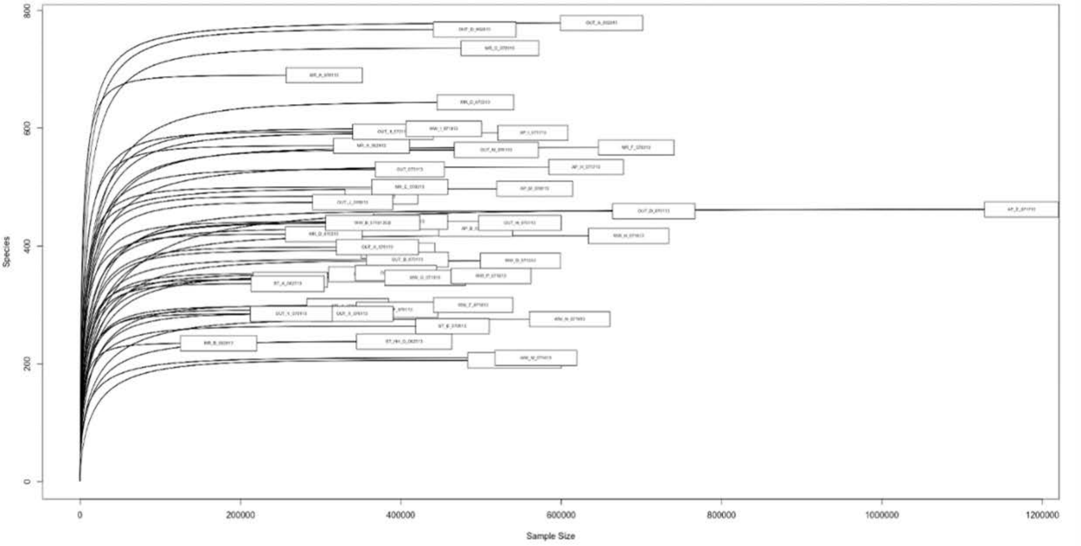

### Supplementary Figure 2

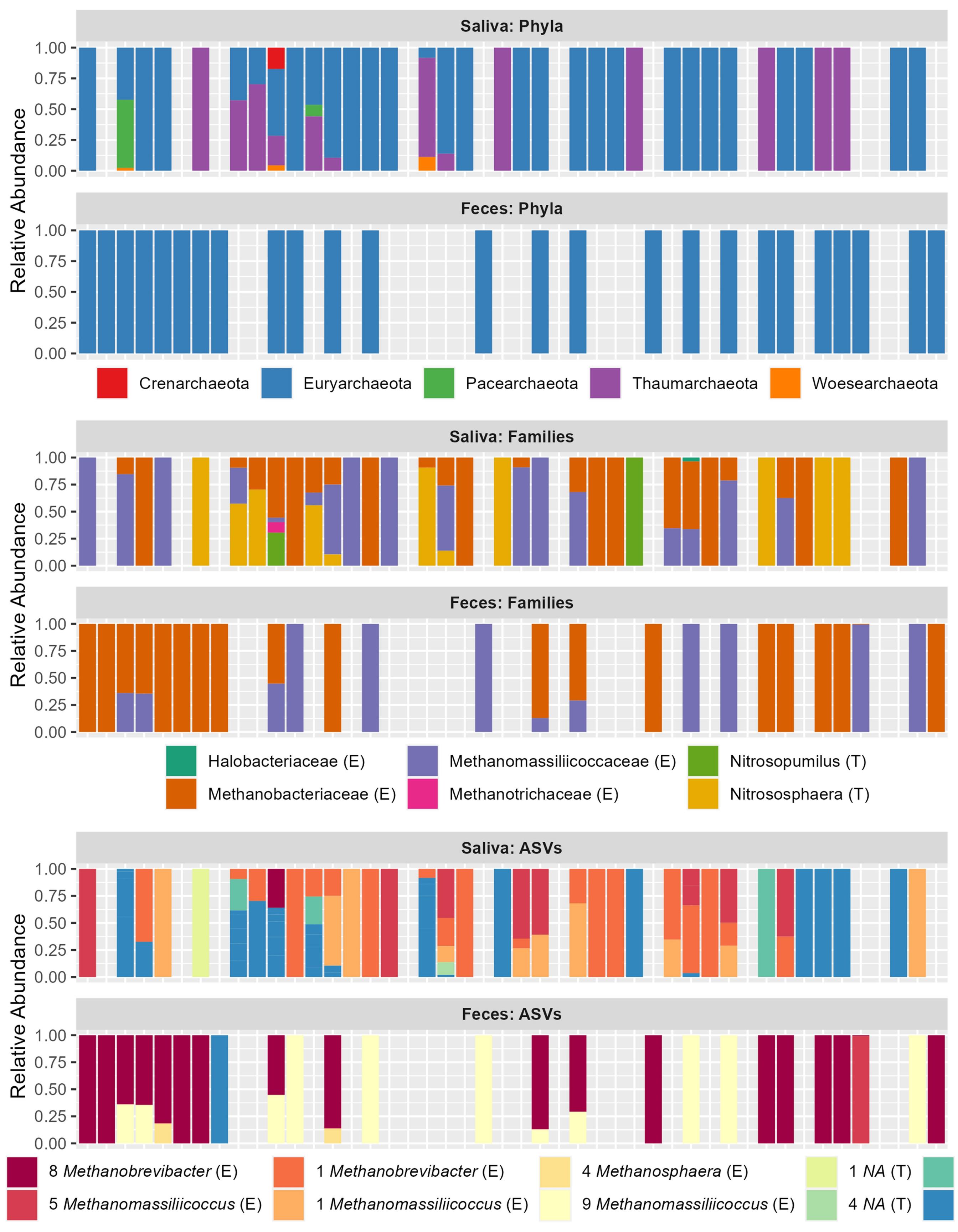
