## Supplementary Table S1 for "Characterizing Bacterial and Archaeal Microbiomes of Singapore’s Urban Long-Tailed Macaques"

Table S1. Mean Alpha Diversity metrics by taxonomic level and region.

| | Oral (N=20) | | | Gut (N=31) | | |
|---|---|---|---|---|---|---|
| | **Richness** | **Evenness** | **Diversity** | **Richness** | **Evenness** | **Diversity** |
| **ASV** | 742.6 | 0.6603288 | 4.324544 | 442.7419 | 0.4926594 | 2.999321 |
| **Family** | 91.8 | 0.558957 | 2.510507 | 62.96774 | 0.4353139 | 1.799877 |
| **Phylum** | 18.95 | 0.5249431 | 1.535187 | 11.3871 | 0.3573008 | 0.8614791 |
